# Specification of the interstitial cell lineage in the fetal mouse ovary requires balanced Notch signaling

**DOI:** 10.64898/2026.04.20.719649

**Authors:** Yu-Ying Chen, Saniya Rattan, Chang Liu, Xin Xu, Humphrey H-C Yao

**Affiliations:** Reproductive Developmental Biology Group, National Institute of Environmental Health Sciences, Research Triangle Park, NC 27709, USA; Epigenetic and RNA Biology Laboratory, National Institute of Environmental Health Sciences, Research Triangle Park, NC 27709, USA

**Keywords:** Ovary, Interstitial cell, Supporting cell, Notch signaling

## Abstract

The ovary contains two major somatic lineages, granulosa cells and interstitial cells, that arise from progenitors within the coelomic epithelium. However, how these two lineages diverge during ovarian development remains unclear. By analyzing joint single-nucleus transcriptomic and chromatin accessibility profiles of murine ovarian cells at the onset of ovary formation, we identified two somatic progenitor populations from the coelomic epithelium distinguished by expression of the nuclear receptors *Nr5a1* and *Nr2f2*. Based on their transcriptomic trajectories, the *Nr5a1*^*+*^ epithelial cells preferentially transitioned toward the granulosa lineage whereas the *Nr2f2*^*+*^ epithelial cells differentiated into mesenchymal populations. This lineage relationship was supported by *Nr2f2* lineage tracing experiments that fetal *Nr2f2*^*+*^ progenitors contribute to ovarian interstitial cells postnatally. To define the molecular features underlying this divergence, we performed differential gene expression and chromatin accessibility analyses and found that *Nr2f2*^*+*^ epithelial cells, but not *Nr5a1*^*+*^ cells, were enriched for Notch pathway components and Notch effector motifs. Consistently, lineage tracing of Notch-responsive cells marked *Nr2f2*^*+*^ interstitial cells in postnatal ovaries, whereas ectopic Notch activation in *Nr5a1*^*+*^ cells promoted expansion of the interstitial population accompanied by reduced granulosa cells. By integrating motif analysis with accessible chromatin-gene linkage, we also identified downstream targets regulated by Notch effectors in *Nr2f2*^*+*^ cells, which showed concordant changes upon ectopic Notch activation. These findings demonstrate that somatic cell fate is established early during ovarian development, with active Notch signaling specifying the interstitial lineage and a balanced Notch activity required for proper somatic lineage establishment.

**Significance Statement:** Proper differentiation of somatic cell types in the fetal ovary lays the foundation for future ovarian function in adulthood. Understanding how each cell type is formed is essential for developing methods to intervene in ovarian diseases caused by cellular dysfunction. Given that common somatic progenitors give rise to both supporting and interstitial lineages, a main unanswered question is how these two lineages diverge apart from each other during ovarian development. By integrating joint single-nucleus transcriptomic and chromatin accessibility assays with lineage tracing, single cell RNA-sequencing, and mouse genetic models, we demonstrate the role of Notch signaling in specifying the interstitial lineage and separating it from the supporting cell fate.

## Introduction

The mammalian ovary develops from the gonadal primordium, which can arise from the coelomic epithelium along the surface of the mesonephros around embryonic day (E) 10 in mice (1, 2). The coelomic epithelium proliferates, thickens, and contributes to two major ovarian somatic lineages: supporting/granulosa cells and interstitial cells (2, 3). The supporting lineage begins with pre-supporting cells positive of *Runx1* expression (4), which subsequently differentiate into pre-granulosa cells positive of *Foxl2* (4). These pre-granulosa cells surround the primordial germ cells that migrate from the hindgut into the gonad (5, 6) forming germ cell cysts (7). These cysts eventually break down into primordial follicles with individual germ cells encapsulated by pre-granulosa cells perinatally (8). Primordial follicles remain non-growing until activated, after which pre-granulosa cells differentiate into granulosa cells to support oocyte growth and maturation (9, 10). In addition, pre-granulosa cells continue to arise from the ovarian epithelium throughout embryonic development and contribute to the second wave of follicle formation (11, 12).

The interstitial lineage comprises both non-steroidogenic stromal cells and steroidogenic theca cells, which reside between follicles, outside the basement membrane that separates granulosa cells from the interstitium/mesenchyme (13). Unlike the epithelial-like pre-granulosa and granulosa cells, interstitial cells are mesenchymal in nature. They provide structural support to the ovary and contribute to extracellular matrix production (13). In the case of theca cells, in response to hormone stimulation they produce androgens, which is subsequently converted to estrogen by granulosa cells (13). Our group demonstrated, using mouse genetic models, that interstitial cells arise from *Wt1*^*+*^ somatic progenitors and *Gli1*^*+*^ mesonephric progenitors recruited during fetal development (14). As *Wt1*^*+*^ progenitors give rise not only to interstitial but also to supporting lineages (14, 15), an unresolved question is how interstitial progenitors diverge from the supporting cell lineage during ovarian development.

Notch signaling is a highly conserved pathway essential for numerous developmental processes, including cell fate specification (reviewed in (16)). In mammals, Notch signaling is activated when ligands of the Jagged or Delta-like families bind to one of four Notch receptors (Notch1-4). Ligand-receptor interaction triggers proteolytic cleavage of the receptor, releasing the Notch intracellular domain to be translocated to the nucleus and forms a transcriptional complex with the DNA-binding protein RBPJ and co-activators of the MAML family. This complex regulates the expression of downstream target genes, including members of the *Hes* and *Hey* families of transcription factors. Conversely, Notch signaling is negatively regulated by NUMB, which promotes receptor degradation and suppresses pathway activation (16). Notch signaling plays critical roles during ovarian development, including follicle formation, follicle growth, and meiotic maturation (17). Loss of Notch ligand *Jag1* in germ cells and receptor *Notch2* in granulosa cells result in follicles containing multiple germ cells, indicating disrupted follicle assembly (18). In addition, global deletion of *Hes1* decreases germ cell numbers while increasing pre-granulosa cells, whereas Notch activation in pre-granulosa cells increases oocyte numbers and reduces pre-granulosa cell abundance (19). Furthermore, *Numb* has been associated with asymmetric division of the coelomic epithelium in the fetal mouse ovary, and its global deletion prior to ovarian formation leads to increased expression of undifferentiated cell markers and a reduction in supporting cells by E13.5 (20). These findings underscore the requirement of regulated Notch activity in somatic cell differentiation and ovarian function.

Given the shared progenitor origin of ovarian somatic lineages, we sought to determine whether differential Notch signaling activity contributes to the divergence of supporting and interstitial cell fates during early ovarian development. We hypothesized that active Notch signaling in epithelial progenitors promotes interstitial lineage specification, whereas reduced Notch activity favors supporting cell differentiation during early gonad development. To address this, we combined single-nucleus transcriptomic and chromatin accessibility profiling at the onset of ovary formation with lineage tracing, single-cell RNA sequencing, and mouse genetic models to investigate how Notch signaling influences early ovarian somatic lineage specification.

## Results

### Identification of two coelomic epithelial populations with differential expression of Notch signaling components

To understand how distinct somatic lineages are established during ovary formation, we analyzed our published joint single-nucleus transcriptomics (RNA) and chromatin accessibility (ATAC) (10x Multiome) dataset (21), and focused on charactering cell type composition, gene expression, and chromatin accessibility of XX murine gonads at embryonic day (E) 11.5, a timepoint that marks the beginning stage of fetal ovary development. A total of 12,335 nuclei were profiled and assigned into 11 major cell populations upon clustering analysis based on RNA expression. They include coelomic epithelial (Epi) positive of *Upk3b* and *Aldh1a*2 (22, 23), mesenchymal (ME) positive for *Ptn* and *Nr2f2* (24), pre-supporting (PSC) positive for *Runx1* and *Wnt4* (4), supporting-like (SLC) positive of *Pax8* (25), germ, endothelial, and immune cells (**Fig. 1A, i & Fig. S1A**). On the other hand, ATAC-based clustering identified 9 chromatin states (**Fig. 1A, ii**), with major cell types corresponded to their respective ATAC clusters (**Fig. 1A, ii, iii & Fig. 1B**). Integrated RNA and ATAC profiles produced well resolved clusters when colored by RNA-derived cell identity, indicating concordance between transcriptional and chromatin accessibility profiles (**Fig. 1A, iv**).

**Figure 1.**
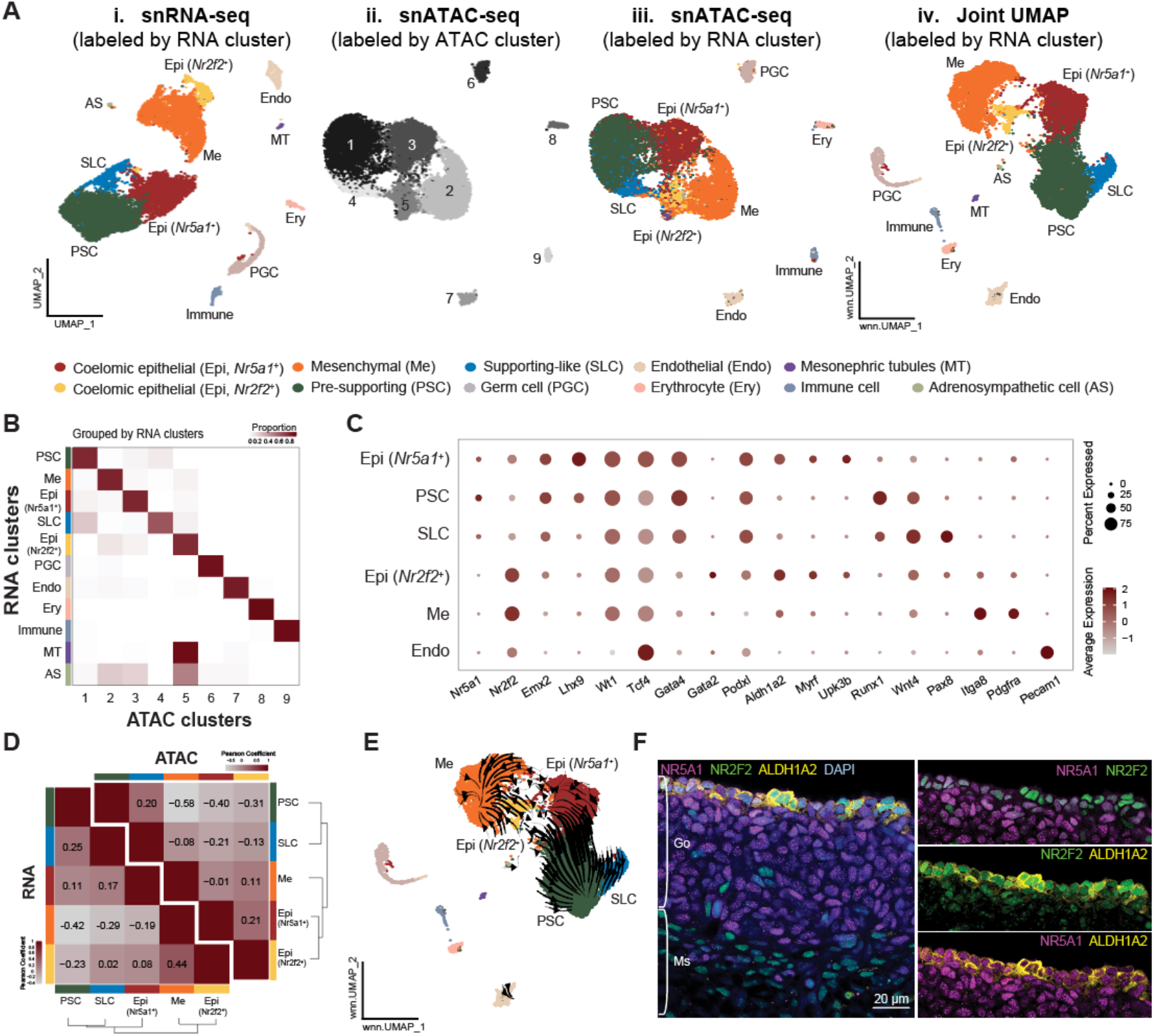
Joint snRNA- and ATAC-seq of murine fetal ovaries revealed two coelomic epithelial populations with differential expression of *Nr5a1* and *Nr2f2*. **(A)** UMAP embeddings of snRNA-seq (i), snATAC-seq (ii-iii), and joint RNA and ATAC (iv) data from E11.5 XX gonadal cells. Cells are color coded by unbiased clustering derived from RNA (i, iii, iv) or ATAC data (ii). Annotations for the snRNA-seq clusters include: coelomic epithelial (Epi, *Nr5a1*^*+*^ or *Nr2f2*^*+*^), mesenchymal (Me), pre-supporting (PSC), supporting-like (SLC), primordial germ cells (PGC), endothelial (Endo), erythrocyte (Ery), Mesonephric tubule (MT), immune, and adrenosympathetic (AS) cells. **(B)** Heatmap of cluster distribution of cells based on ATAC clustering (x-axis) and RNA-clustering (y-axis), as grouped by individual RNA cluster (the sum of each row is 1). **(C)** Marker gene expression for cell type annotation based on RNA clustering. **(D)** Hierarchical clustering with Pearson correlation coefficient of top variable genes (bottom left) and top variable chromatin regions (top right) across major somatic clusters. **(E)** Velocity stream computed using MultiVelo overlaid on joint RNA and ATAC UMAP of E11.5 XX gonadal cells. **(F)** Immunostaining of E11.5 wildtype XX gonads (Go) and mesonephros (Ms) with NR5A1 (magenta), NR2F2 (green), ALDH1A2 (yellow), and DAPI (blue) staining. Scale bar= 20µm.

Within the somatic cell clusters, we identified two distinct coelomic epithelial populations. Although both expressed canonical epithelial markers such as *Aldh1a2* and *Podxl* (24), one epithelial cluster was enriched for the nuclear receptor *Nr5a1* (or Steroidogenic factor 1, *Sf1*), whereas the other preferentially expressed *Nr2f2* (or *Coup-tfII*; **Fig. 1C**). Pearson correlation and hierarchical clustering using the RNA data revealed that the *Nr5a1*^+^epithelial cluster was transcriptionally more similar to pre-supporting cells and supporting-like cells, while the *Nr2f2*^+^epithelial population aligned more closely with mesenchymal cells (**Fig. 1D**, bottom left). In contrast, ATAC-based correlation and clustering placed the two epithelial clusters closer to each other than other lineages, followed by mesenchymal and supporting populations (**Fig. 1D**, top right). This observation implies that lineage transitions/separation may be driven more by cluster-specific transcription factor program than by a broad shift in chromatin landscape. To investigate potential lineage trajectories, we applied MultiVelo, an RNA velocity inference tool that integrates both RNA splicing dynamics and chromatin accessibility changes for cell fate prediction (26) on the joint UMAP (**Fig. 1A, iv**). Using this tool, we observed velocity streams originating from *Nr5a1*^+^epithelial cells toward pre-supporting cells, consistent with a developmental path leading to pre-granulosa cells (**Fig. 1E**). In contrast, a distinct velocity stream originated from *Nr2f2*^+^ epithelial cells and progressed toward mesenchymal populations, suggesting the potential contribution of *Nr2f2*^+^ epithelial cells to the ovarian interstitial lineage (**Fig. 1E**). We also note that supporting-like cells (SLC) exhibited a distinct velocity stream (**Fig. 1E**), aligning with prior reports that despite their close association with the coelomic epithelium at E10.5 (25), the SLCs appear transcriptionally and chromatin-wise distinct by E11.5 and cluster further from epithelial progenitors (**Fig. 1D**).

Consistent with these observations, whole-mount immunostaining of E11.5 female gonads showed inverse expression pattern of NR5A1 and NR2F2 proteins in the coelomic epithelium (**Fig. 1F, Fig. S1B**). Together, these observations reveal a model in which *Nr5a1*^+^epithelial cells give rise to supporting-cell lineages, whereas *Nr2f2*^+^epithelial cells contribute to the interstitial lineages.

### Notch signaling and Notch-related transcription factor motifs are enriched among *Nr2f2*^+^ epithelial cells

To identify molecular features distinguishing the *Nr5a1*^+^and *Nr2f2*^+^epithelial populations, we analyzed both differentially expressed genes (DEGs) and differential accessible chromatin peaks (DAPs) between these two populations at E11.5. At the transcriptomic level, *Nr2f2*^+^epithelial cells displayed 729 DEGs (Log2FC >1, adjusted p-value < 0.05) with enriched molecular signatures spanning epithelial mesenchymal transition, Hedgehog signaling, angiogenesis, and Notch signaling (**Fig. 2A, B**, top). In contrast, *Nr5a1*^+^epithelial cells showed 359 DEGs with pathways enrichment primarily associated with hormone-response signaling (**Fig. 2A, B**, bottom). Notably, several Notch pathway components, including *Hes1, Maml2, Notch3, Jag1*, were enriched in *Nr2f2*^+^epithelial and mesenchymal cells, whereas the Notch antagonist *Numb* was highly expressed in *Nr5a1*^+^epithelial and supporting cells, including both pre-supporting and supporting-like cells (**Fig. 2C, Fig. S2A**). Endothelial cells also exhibited strong expression of Notch effectors, consistent with established roles of Notch signaling in vascular development (27) (**Fig. 2C**). These findings suggest that differential Notch signaling activity may contribute to fate divergence between supporting and interstitial lineages.

**Figure 2.**
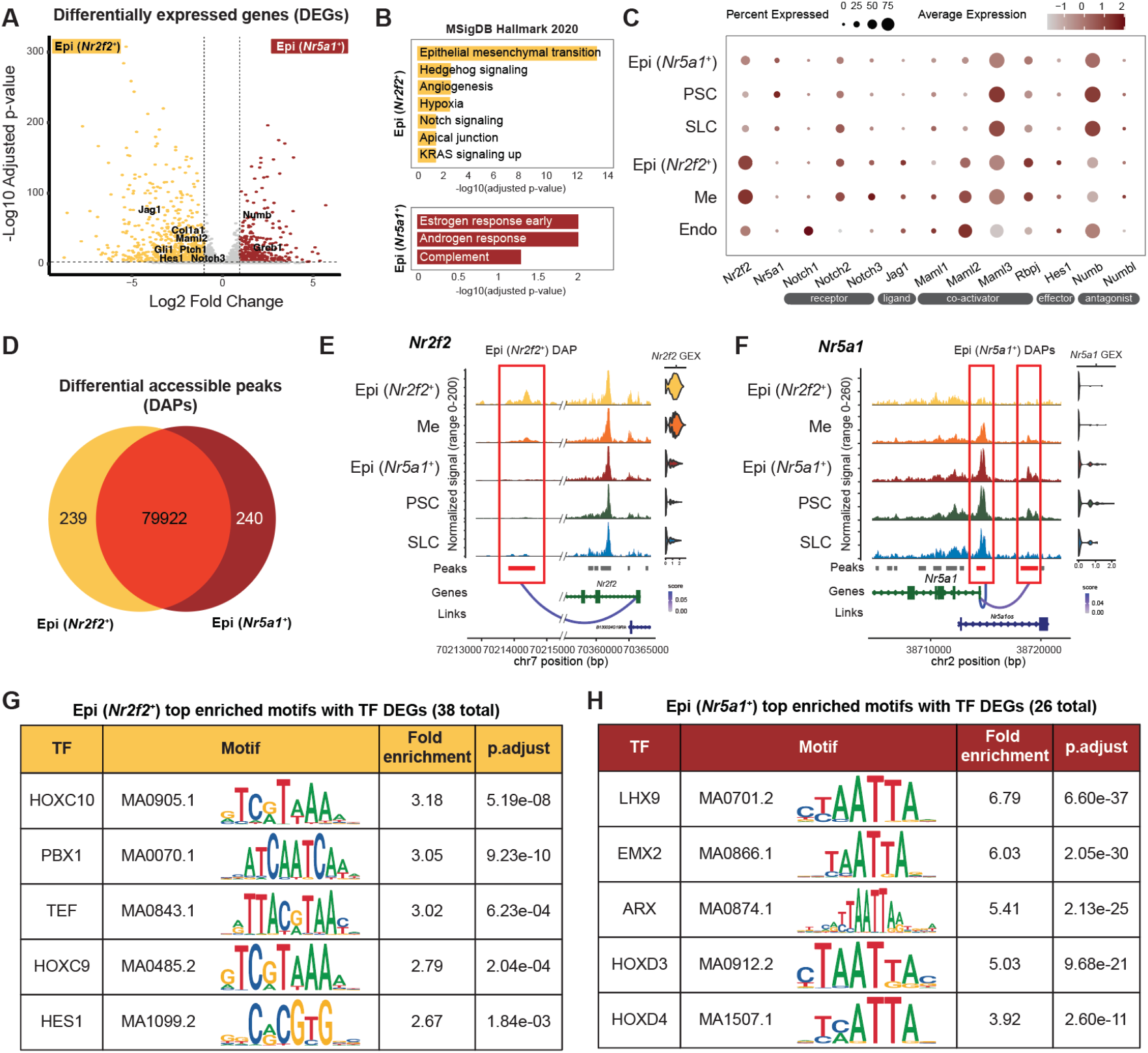
Notch signaling and Notch-related transcription factor motifs are enriched among *Nr2f2*^*+*^ epithelial-enriched accessible chromatin regions. **(A)** Volcano plot of differentially expressed genes (DEGs) between Epi, *Nr2f2*^*+*^ (yellow) versus Epi, *Nr5a1*^*+*^ (red) cells. **(B)** Pathway analysis using MSigDB Hallmark 2020 database showing enriched pathways among DEGs upregulated in Epi, *Nr2f2*^*+*^ (yellow) versus Epi, *Nr5a1*^*+*^ (red) cells. **(C)** Dot plot of the expression of various Notch signaling components across major somatic cell types. **(D)** Venn diagram comparing accessible chromatin peaks between *Nr2f2*^*+*^ epithelial and *Nr5a1*^*+*^ cells. **(E-F)** Coverage plots illustrating *Nr2f2*^*+*^ epithelial (E) and *Nr5a1*^*+*^ epithelial-enriched (F) DAPs and their associated peak-gene linkages. Normalized chromatin accessibility is shown on the left and gene expression (GEX) on the right. Red boxes and red bars highlight DAPs that are positively linked to their associated gene, whereas gray bars denote peaks that were detected but not differentially accessible. “Score” represents the correlation coefficient between peak accessibility and gene expression. **(G-H)** Top five significantly enriched transcription factor motifs identified in *Nr2f2*^*+*^ (G) or *Nr5a1*^*+*^ (H) epithelial-specific DAPs, ranked by fold enrichment.

We next compared the chromatin profile between *Nr2f2*^+^ and *Nr5a1*^+^ epithelial cells at E11.5. Differential analysis revealed 239 *Nr2f2*^+^ epithelial-enriched and 240 *Nr5a1*^+^ epithelial-enriched DAPs (Log2FC >0, adjusted p-value < 0.05) among over 80,000 accessible chromatin regions profiled (**Fig. 2D**), consistent with the overall similarity in chromatin profile between the two clusters (**Fig. 1D**). These DAPs mapped to genomic regions largely to distal intergenic (46 - 49.6%), intronic regions (33.3 - 38.9%), and about 10% in the promoter regions (**Fig. S2B**). To assess whether differential chromatin accessibility was associated with transcriptional output, we performed peak-gene linkage analysis by computing correlations between chromatin peak accessibility and gene expression within the linkage distance of transcription start site (TSS) ± 500Kb. Among the 239 *Nr2f2*^+^ epithelial-enriched DAPs, 16.0% were linked to *Nr2f2*^+^ epithelial-enriched DEGs, corresponding to 42 unique genes including *Nr2f2* and *Col1a1*. In the case of *Nr5a1*^+^ epithelial-enriched DAPs, 65 unique genes were linked to 24.1% of *Nr5a1*^+^-enriched DAPs, including *Nr5a1* and *Amhr2* (**Fig. S2C, SI Data**). Notably, DEGs without any associated differential chromatin regions accounted for the majority of total DEGs in both populations, suggesting that expression of these genes may be regulated through different transcription factor binding in pre-accessible chromatin regions not differential between the two epithelial populations or *via* distal regulatory elements beyond the defined linkage distance.

An example of DAP enriched in *Nr2f2*^+^ epithelial cells was a peak located downstream of *Nr2f2* gene body itself, which was highly accessible in *Nr2f2*^*+*^epithelial cells, reduced in mesenchymal cells, and absent in *Nr5a1*^+^ epithelial and supporting lineages (**Fig. 2E**).

This peak was positively linked to *Nr2f2* expression (**Fig. 2E**). Conversely, *Nr5a1*-enriched DAPs included both the *Nr5a1* promoter peak and an upstream peak from its gene body. The *Nr5a1* promoter peak was accessible in all lineages except *Nr2f2*^*+*^epithelial cells, whereas the upstream peak was accessible in *Nr5a1*^*+*^epithelial and pre-supporting cells, and showed decreased accessibility in mesenchymal, supporting-like, and *Nr2f2*^*+*^epithelial cells (**Fig. 2F**).

Given that a subset of DAPs showed positive correlations with gene expression, we sought to identify potential transcription factors (TFs) that may regulate lineage-specific gene expression. Motif enrichment analysis within the DAPs, intersected with DEG lists, identified candidate TFs whose motifs were enriched and whose transcripts were differentially expressed. In *Nr2f2*^*+*^epithelial cells, 38 TFs met these criteria, including members of the HOX family, PBX1, TEF, and Notch effector HES1 (**Fig. 2G**). Another Notch co-activator RBPJ was also an enriched motif, however, it was not a DEG between the two epithelial populations (**SI Data**). In *Nr5a1*^*+*^epithelial cells, 26 TFs showed both motif enrichment and differential expression, including LHX9, EMX2, ARX (28), and several HOX family TFs, all characterized by variants of the core “TAATTA” homeobox motif (**Fig. 2H**). Notably, LHX9 and EMX2 have been proposed as regulators of pre-granulosa cell differentiation (29). NR5A1, LEF1, and GATA4 were also among the enriched motifs in *Nr5a1*^*+*^epithelial cells (**SI Data**).

### Notch-responsive and *Nr2f2*-positive cells in the fetal ovary give rise to the interstitial cell population in postnatal ovaries

To determine whether ovarian interstitial cells derive from Notch-responsive and/or *Nr2f2*^+^progenitors, we performed tamoxifen-induced lineage tracing experiments by crossing inducible Notch effector *Hes1*^*CreERT2*^ or *Nr2f2*^*iCreERT2*^ mice to *ROSA*^*tdTomato*^ reporter allele separately. Upon tamoxifen administration, *Hes1*^+^ (Notch-responsive) or *Nr2f2*^+^cells in the embryos (*Hes1*^+/*CreERT2*^; *ROSA*^*+/tdTomato*^ or *Nr2f2*^*+/iCreERT2*^; *ROSA*^*+/tdTomato*^) were permanently labeled with tdTomato fluorescent proteins. Tamoxifen was given to pregnant dams as a single injection at E10.5, E11.5, or E13.5, and ovaries from F1 pups were collected at E11.5, E16.5, and postnatal day (P) 21 (**Fig. 3A**).

**Figure 3.**
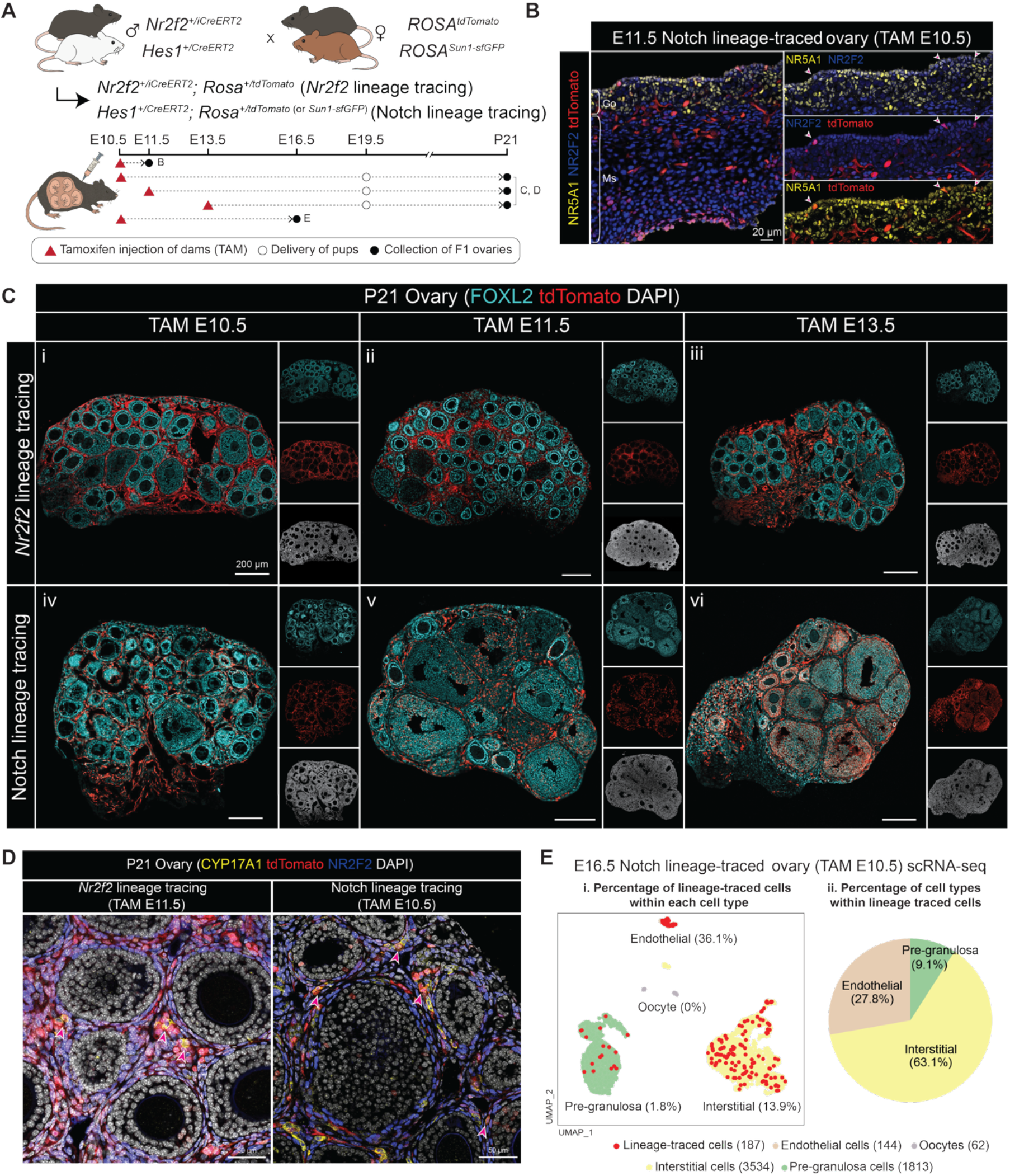
Notch-responsive and Nr2f2-positive cells at E10.5-E11.5 give rise to the interstitial cell population in postnatal ovaries. **(A)** Schematic of the genetic strategy for Notch-responsive and *Nr2f2*^+^progenitor lineage tracing models. Tamoxifen injection (TAM) is labeled with red triangles, ovary collection is labeled with black circles. White circles indicate terminal C-section and fostering for postnatal collections. **(B)** Immunostaining of E11.5 *Hes1*^+/CreERT2^; *ROSA*^*+/tdTomato*^ gonad (Go) and mesonephros (Ms) lineage-traced at E10.5, stained with NR5A1 (yellow), NR2F2 (blue), and tdTomato (red). Arrows mark lineage-traced cells in the coelomic epithelium of the gonadal side. Scale bar= 20µm. **(C)** Immunostaining of P21 *Nr2f2*^*+/iCreERT2*^; *ROSA*^*+/tdTomato*^ (*Nr2f2* lineage tracing: i-iii) and *Hes1*^+/CreERT2^; *ROSA*^*+/tdTomato*^ (Notch lineage tracing: iv-vi) ovaries lineage-traced at E10.5 (i, iv), E11.5 (ii, v), or E13.5 (iii, vi), with FOXL2 (cyan), tdTomato (red), and DAPI (gray) staining. Left panels show merged cyan and red channels; right panels show single channels. Scale bar= 200µm. **(D)** Immunostaining of P21 *Nr2f2*^+/CreERT2^; *ROSA*^*+/tdTomato*^ ovaries lineage-traced at E11.5 (left) and P21 *Hes1*^*+/iCreERT2*^; *ROSA*^*+/tdTomato*^ ovaries lineage-traced at E10.5 (right), marked with CYP17A1 (yellow), dtTomato (red), NR2F2 (blue), and DAPI (white). Red arrows mark lineage-trace cells expressing CYP17A1. Scale bar= 50µm. **(E)** UMAP of single-cell RNA-seq from E16.5 *Hes1*^+/*CreERT2*^; *ROSA*^*+/Sun1-sfGFP*^ ovaries lineage-labeled at E10.5. Left panel shows the percentage of lineage-traced cells (*sfGfp*^*+*^) within each annotated cell type, with total cell number per cell type indicated (i). Right panel shows the proportional cell type composition within the lineage-traced population (ii).

To assess the identity and distribution of Notch-responsive cells at the onset of ovarian formation, fetal ovaries from *Hes1*^+/*CreERT2*^; *ROSA*^*+/tdTomato*^ embryos were collected at E11.5, 24 hours after tamoxifen administration. The majority of lineage-traced cells, as labeled by tdTomato staining, were positive for NR2F2 and negative of NR5A1. This finding aligns with our model that Notch signaling is active in the *Nr2f2*^+^ cells in the undifferentiated ovary (**Fig. 3B**). Lineage-traced cells were detected in both the gonad and mesonephros and were distributed among endothelial, mesenchymal, and the coelomic epithelium (**Fig. S3A**).

We next examined what *Nr2f2*^+^and *Hes1*^+^/Notch-responsive cells at E10.5 became in P21 ovaries. In the *Nr2f2*^*iCreERT2*^ lineage tracing model, tamoxifen administration at E10.5 resulted in labeling of predominantly interstitial cells and a small number of granulosa cells in P21 ovaries (**Fig. 3C, i**). When tamoxifen was given later at E11.5 or E13.5, nearly all labeled cells were confined to the interstitial compartment, including CYP17A1^+^ steroidogenic theca cells, with minimal labeling of supporting cells (**Fig. 3C, ii, iii, Fig. 3D**). In the *Hes1*^+/CreERT2^ lineage tracing model, we found that Notch-responsive cells at E10.5 became interstitial cells postnatally, including CYP17A1^+^ steroidogenic theca cells, and a small number of granulosa cells (**Fig. 3C, iv, Fig. 3D, Fig. S3B**). However, induction at E11.5 and E13.5 resulted in a progressively greater contribution of lineage-traced cells to the granulosa population, indicating that granulosa precursors exhibit Notch activity during this time (**Fig. 3C, v, vi**). These results suggest that progenitor cells positive of both *Nr2f2* expression and Notch activity around E10.5-E11.5 contribute to the interstitial lineage.

To more precisely characterize the identity of lineage-traced cells, we performed single-cell RNA sequencing on E16.5 *Hes1*^+/*CreERT2*^; *ROSA*^*+/Sun1-sfGFP*^ ovaries that were tamoxifen induced at E10.5. The *ROSA*^*+/Sun1-sfGFP*^ reporter was used in place of the *ROSA*^*+/tdTomato*^ reporter because it produced lower background signals during transcript alignment. A total of 5,553 cells were analyzed and clustered into oocyte, pre-granulosa, interstitial, and endothelial populations (**Fig. S4A**). Among these, 187 *Gfp*^+^ lineage-traced cells were identified, constituting about 36.1% of all endothelial cells, 13.9% of interstitial cells, 1.8% of pre-granulosa cells, with none of the oocytes being labeled (**Fig. 3E, i**). Within the *Gfp*^+^ traced population, 63.1% were interstitial, 27.8% endothelial, and less than 10% supporting cells (**Fig. 3E, ii**). Among the Notch-related genes, *Notch1, Notch3, Maml2, Heyl, Wnt5a* were DEGs of interstitial cells; however, *Hes1* and *Rbpj* expression were also observed in supporting cells, indicating that Notch is active also in the supporting cells at E16.5 as previously suggested (18, 19) (**Fig. S4B**). Overall, these data suggest that, at E10.5, *Hes1*^+^/Notch-responsive progenitors at E10.5 predominantly adopt interstitial fates.

### Ectopic Notch activation in *Nr5a1*^*+*^ cells increases interstitial-to-supporting cell ratio, upregulates interstitial gene expression, and disrupts ovarian morphology

Based on our finding that Notch-responsive and/or *Nr2f2*^*+*^ cells mark interstitial progenitors (**Fig. 3**), we hypothesized that active Notch signaling in *Nr2f2*^*+*^ cells at the onset of ovary formation specifies the interstitial lineage fate, whereas repressed Notch signaling in the *Nr5a1*^*+*^ cells permits supporting cell differentiation. To test this, we ectopically activated the Notch pathway in *Nr5a1*^*+*^ cells by crossing Tg(Nr5a1-cre) mice with *ROSA*^*Notch*^ mice, generating embryos expressing a constitutively active Notch intracellular domain in the *Nr5a1*^*+*^ cells. Fetal ovaries from Notch-induced (Nr5a1-cre+; *ROSA*^*+/Notch*^) and control (Nr5a1-cre+; *ROSA*^+/+^) littermates were collected at E13.5, E15.5, and E18.5 for analysis (**Fig. 4A**).

**Figure 4.**
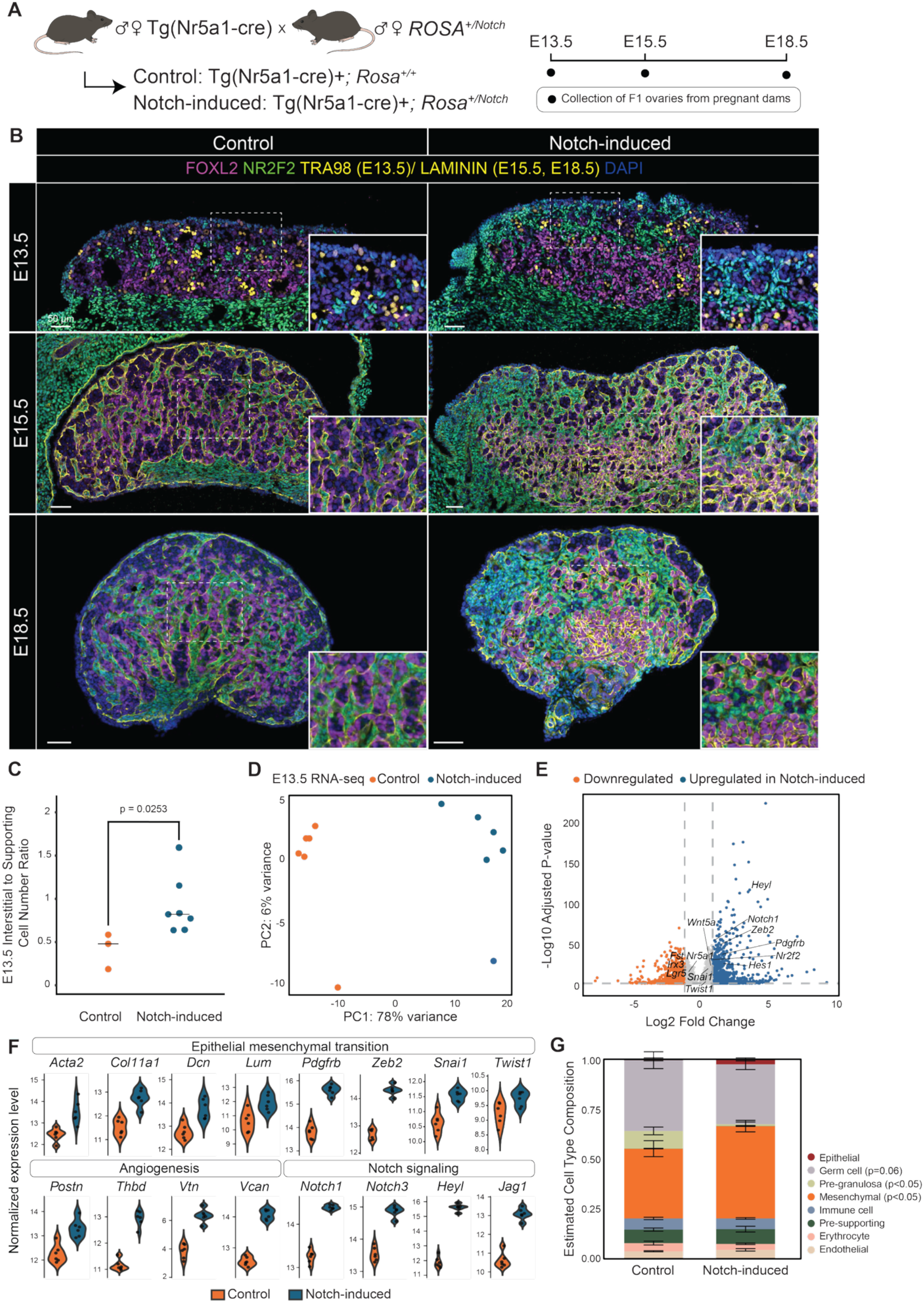
Ectopic Notch activation in *Nr5a1*^*+*^ cells increases interstitial-to-supporting cell ratio, upregulates interstitial gene programs, and disrupts ovarian morphology. **(A)** Schematic of the genetic strategy for ectopic Notch activation. **(B)** Immunofluorescence of control and Notch-induced ovaries at E13.5 (top), E15.5 (middle), and E18.5 (bottom) stained with FOXL2 (magenta), NR2F2 (green), TRA98 (yellow, E13.5) or LAMININ (yellow, E15.5–E18.5), and DAPI (blue). Scale bar= 50µm. **(C)** Quantification of the NR2F2^+^(interstitial) to FOXL2^+^(supporting pre-granulosa) cell ratio at E13.5 in Notch-induced versus control ovaries. **(D)** Principal component analysis (PCA) of bulk RNA-seq from E13.5 control and Notch-induced ovaries. **(E)** Volcano plot of DEGs in E13.5 Notch-induced versus control ovaries. **(F)** Normalized expression of representative genes within MSigDB Hallmark pathways enriched among DEGs upregulated in Notch-induced ovaries. **(G)** Single-cell deconvolution of bulk RNA-seq data using E13.5 snRNA-seq as reference.

At E13.5, control ovaries contained on average half as many NR2F2^+^ interstitial cells as FOXL2^+^ supporting pre-granulosa cells. In contrast, Notch-induced ovaries exhibited a significant increase in interstitial relative to supporting cells, with interstitial cell numbers equal to or exceeding those of supporting cells (**Fig. 4B**, top, **Fig. 4C**). Moreover, NR2F2^+^ cells were frequently observed in clusters near the ovarian epithelium (**Fig. 4B**, top). The presence of germ cells, supporting cells, and interstitial cells in Notch-induced ovaries indicates that ectopic Notch activation in *Nr5a1*^*+*^ cells does not prevent initial ovary formation. By E15.5, morphological differences between the two genotypes became more apparent. In control ovaries, laminin staining delineated well-organized germ cell cysts in which oocyte clusters were surrounded by pre-granulosa cells. In contrast, Notch-induced ovaries displayed disrupted cyst organization, characterized by smaller, irregular cysts with increased accumulation of interstitial cells between them (**Fig. 4B**, middle). By E18.5, ovarian organization was further compromised in Notch-induced ovaries with reduced oocyte numbers and disorganized pre-granulosa cells that failed to properly surround oocytes (**Fig. 4B**, bottom, **Fig. S5**).

To characterize the molecular changes of ectopic Notch activation, we performed bulk RNA-sequencing on E13.5 control and Notch-induced ovaries. Principal component analysis (PCA) showed clear separation between control and Notch-induced samples along PC1 with 78% variance, indicating a strong genotype-driven transcriptional shift (**Fig. 4D**). Differential gene expression analysis identified a comparable number of upregulated and downregulated DEGs following Notch activation (890 downregulated and 885 upregulated in Notch-induced ovaries, Log2FC > 1.5, adjusted p-value < 0.05); although genes upregulated in Notch-induced ovaries exhibited greater statistical significance (**Fig. 4E**). Among DEGs upregualted in Notch-induced ovaries, genes in the pathways including epithelial-to-mesenchymal transition (e.g. *Col11a1, Dcn, Pdgfrb*, and *Snai1*), angiogenesis (e.g. *Postn*), and Notch signaling (e.g. *Notch1, Notch3, Heyl, Jag1*) were significantly enriched (**Fig. 4F**), whereas the only downregulated pathway was heme metabolism. To assess whether ectopic Notch activation alters cell type composition, we performed deconvolution of the bulk RNA-seq data using an E13.5 single-nucleus RNA-seq dataset (21) as our reference data. This analysis revealed a significant increase in mesenchymal cell abundance and a concomitant decrease in pre-granulosa cells in Notch-induced ovaries. A trend toward reduced germ cell abundance was also observed (adjusted *p* = 0.06) (**Fig. 4G**). Together, these findings indicate that ectopic Notch activation in *Nr5a1*^*+*^ cells drives a shift toward a mesenchymal gene expression program, and a balanced Notch signaling is essential for establishing proper somatic cell composition during ovarian development.

### Interstitial gene expression is linked to Notch-related transcription factor motifs

To delineate how Notch signaling regulates interstitial-related gene expression in the fetal ovary, we performed motif analysis using the joint single-nucleus RNA and ATAC dataset. We first identified differential accessible chromatin peaks (DAPs) between *Nr2f2*^+^ and *Nr2f2*^-^ cells and used these peaks for transcription factor (TF) motif enrichment analysis (**Fig. 5A, i**). We next performed motif scan within accessible peaks of *Nr2f2*^+^ epithelial and mesenchymal cells to determine motif positions (**Fig. 5A, ii**) and linked putative TFs, including HES1, RBPJ, and NR2F2 to their downstream target genes (**Fig. 5A, iii**).

**Figure 5.**
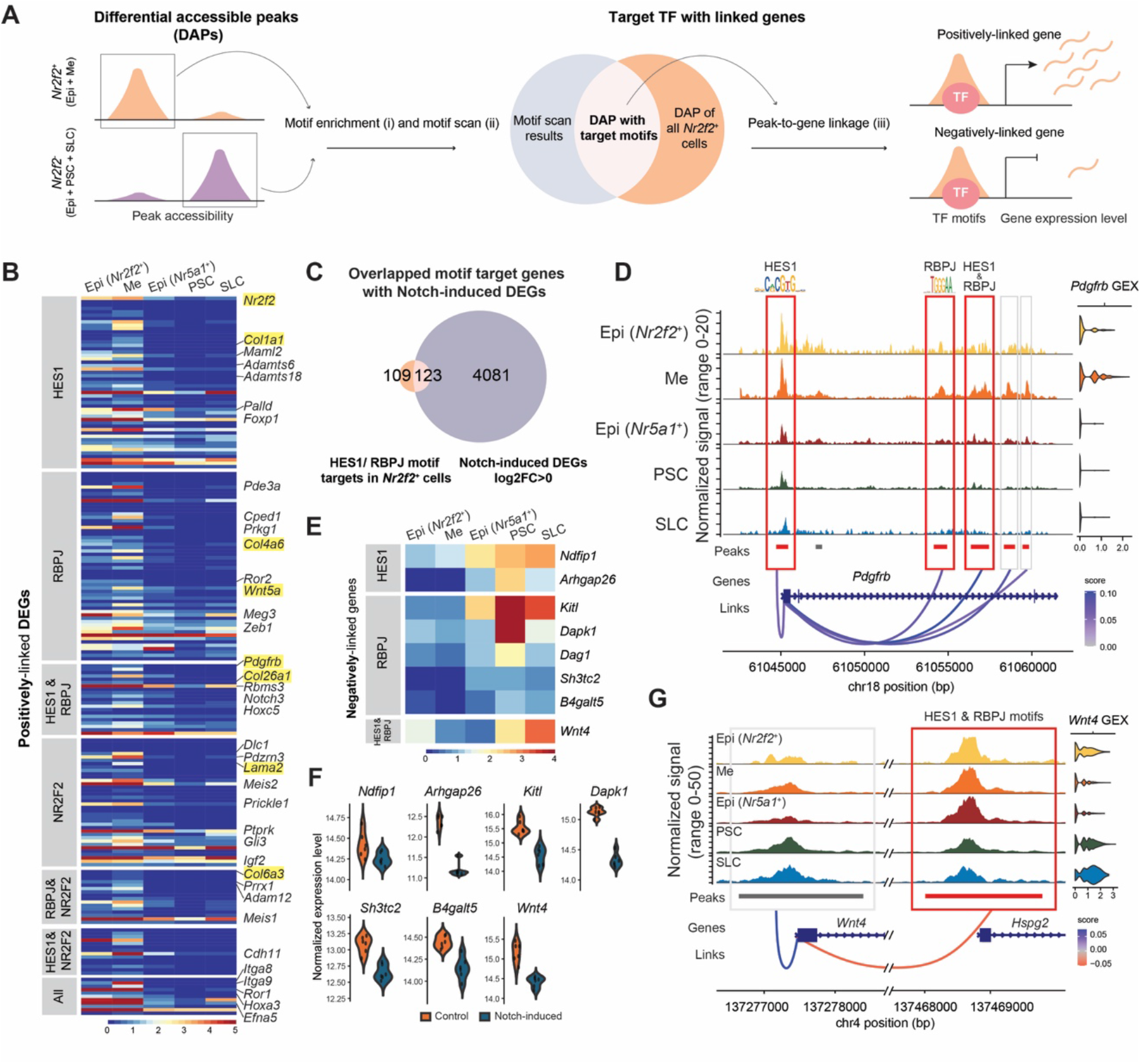
Notch-related transcription factor motifs are linked to interstitial gene expression. **(A)** Workflow of motif enrichment and linkage analysis: (i) Motif enrichment was performed on differentially accessible peaks (DAPs) in *Nr2f2*^*+*^ and *Nr2f2*^*-*^ cells. (ii) Motif scan was performed to determine motif position within DAPs of all *Nr2f2*^*+*^ cells. (iii) Downstream targets of the motifs were identified based on the linkage analysis. **(B)** Heatmap of normalized expression of positively-linked DEGs associated with HES1, RBPJ, and NR2F2 motifs, identified in *Nr2f2*^*+*^ epithelial and mesenchymal cells, across major somatic populations. Linked genes are grouped by inferred regulatory category (grey bars on the left), including HES1-only, RBPJ-only, HES1 & RBPJ, NR2F2-only, RBPJ & NR2F2, HES1 & NR2F2, or jointly regulated by all three motifs. **(C)** Venn diagram showing the overlap between positively linked DEGs from *Nr2f2*^*+*^ DAPs and genes upregulated in E13.5 Notch-induced ovaries (log2FC>0, adjusted p-value <0.05). **(D)** Coverage plot of the *Pdgfrb* locus. Red boxes with red bars denote positively linked DAPs containing mapped HES1 and/or RBPJ motifs; gray boxes with red bars indicate positively linked DAPs lacking these motifs. Gray bars denote peaks detected but not differentially accessible. **(E)** Heatmap showing normalized expression of negatively linked genes associated with HES1 and/or RBPJ motifs. **(F)** Violin plots displaying normalized expression of negatively linked genes identified in (E) in control versus E13.5 Notch-induced ovaries. **(G)** Coverage plot of the *Wnt4* and *Hspg2* loci illustrating an *Nr2f2*^*+*^-enriched DAP that is negatively linked to *Wnt4* expression. The red box with a red bar marks the DAP whose accessibility is negatively associated with *Wnt4* expression and contains both HES1 and RBPJ motifs. The *Wnt4* promoter peak (gray box and gray bar) is not a DAP despite positively-linked to *Wnt4* expression. Epi (*Nr2f2*^*+*^): *Nr2f2*^*+*^ epithelial, ME: mesenchymal, Epi (*Nr5a1*^*+*^): *Nr5a1*^*+*^epithelial, PSC: pre-supporting cells, SLC: supporting-like cells.

Using this workflow, we first identified DAPs between all *Nr2f2*^*+*^ (including *Nr2f2*^*+*^ epithelial and mesenchymal cells) and *Nr2f2*^*-*^ cells (including *Nr5a1*^*+*^ epithelial, pre-supporting, and supporting-like cells). The analysis revealed over 3,000 DAPs in *Nr2f2*^*+*^ cells and over 2,000 in *Nr2f2*^*-*^ cells (**Fig. S6A**). Motif enrichment within *Nr2f2*^*+*^ DAPs revealed significant enrichment for motifs of NR2F2 and Notch-related TFs including RBPJ and HES1 (**SI Data**). Next, we performed motif scan to identify the precise genomic locations of these motifs within the DAPs. To infer potential downstream target genes, we conducted peak-gene linkage analysis, assessing whether accessibility of a given chromatin region positively or negatively correlated with expression of genes with a linkage distance of transcription start site (TSS) ± 500Kb. This approach identified *Nr2f2*^*+*^ cells-enriched DAPs containing RBPJ and/ or HES1 motifs that were positively linked to canonical interstitial markers, including *Pdgfrb, Wnt5a, Nr2f2*, and multiple collagen genes (**Fig. 5B**). DAPs containing the NR2F2 motif were also linked to interstitial gene expression (**Fig. 5B**). When filtered for DEGs enriched in *Nr2f2*^+^ cells, most linked genes showed higher expression in *Nr2f2*^*+*^ epithelial and mesenchymal cells and lower expression in *Nr5a1*^*+*^ epithelial, pre-supporting, and supporting-like cells (**Fig. 5B**). Most predicted target genes were associated with a single TF, with fewer co-regulated by more than one TFs, and very few linked to NR5A1 (**Fig. 5B, Fig. S6B**). To further evaluate whether these predicted targets respond to Notch signaling, we overlapped the candidate target genes of HES1 and RBPJ with DEGs upregulated in E13.5 Notch-induced ovaries (**Fig. 4E**). More than half of these positively linked targets were upregulated following ectopic Notch activation (**Fig. 5C**). One example of such upregulation was *Pdgfrb*, whose expression increased in Notch-induced ovaries and was predicted to be regulated by Notch effectors HES1 and RBPJ. Expression of *Pdgfrb* in *Nr2f2*^*+*^ cells was linked to 5 DAPs, including the promoter peak containing a HES1 motif, one intragenic peak containing an RBPJ motif, and another intragenic peak containing both HES1 and RBPJ motifs (**Fig. 5D**).

We also detected negatively linked genes, defined as genes for which increased chromatin accessibility was associated with decreased gene expression, suggesting potential repressive effects of Notch-related TF binding. Negatively associated genes included *Wnt4, Kitl, Ndfip1*, and *Arhgap26*, which were more highly expressed in *Nr5a1*^*+*^ epithelial, pre-supporting, and supporting-like cells (**Fig. 5E**). Consistently, these genes were downregulated in Notch-induced ovaries (**Fig. 5F**). One example of such a negatively linked region was the *Hspg2* promoter peak containing both HES1 and RBPJ motifs, which showed a negative association with *Wnt4* expression (**Fig. 5G, Fig. S6C**).

Together, these results reveal distinct transcriptional regulatory networks underlying somatic cell lineage separation in the fetal ovary. Notch-related TFs may directly promote interstitial fate specification by activating interstitial-associated gene programs (e.g. *Pdgfrb)* while repressing supporting-associated genes (e.g. *Wnt4*).

## Discussion

In this study, we aimed to address an important unresolved question in ovarian biology: how the two major somatic lineages, supporting granulosa and interstitial cells, establish distinct identities and diverge during development. Previous studies using *ex vivo* fluorescent dye labeling of the coelomic epithelium and BrdU pulse-chase experiments have shown that coelomic epithelial cells overlying the gonad proliferate and migrate inward without apparent lineage restriction toward either supporting or interstitial fates (3, 30). Evidence for the earliest divergence of the two lineages in the fetal ovary comes from mouse genetic lineage-tracing studies using tamoxifen-inducible *Axin2*^*CreERT2*^ model, which marks WNT-responsive cells. When *Axin2*-positive progenitors are labeled at E10.5, they give rise exclusively to supporting cells of first-wave medullary follicles (12). Similarly, lineage tracing using *Foxl2*^*CreERT2*^ (11, 31) and *Lgr5*^*CreERT2*^ (12) at E12.5 and later timepoints label only granulosa cell lineages. In addition, granulosa cells can arise from *Nestin*^+^perivascular cells labeled at E12.5, although labeling at later embryonic stages also gives rise to steroidogenic interstitial cells (32). On the other hand, interstitial cell progenitors have been exclusively traced from *Gli1*^+^mesonephric progenitors at E12.5 (14) and from *Wnt5a*^*+*^ cells at E15.5 (33). In the present study, we demonstrate that progenitors of ovarian interstitial cells can be lineage labeled as early as E10.5, demonstrating that separation of the two somatic lineages occurs at the beginning of ovary formation.

To gain mechanistic insight into how somatic lineage separation occurs, we analyzed our 10x Multiome dataset (21) and identified two coelomic epithelial populations in the E11.5 mouse ovary: one characterized by *Nr2f2* expression and enrichment of Notch effectors, and the other defined by *Nr5a1* expression and enrichment of Notch antagonist *Numb*. Trajectory inference using RNA velocity analysis suggested that *Nr5a1*^*+*^ epithelial cells preferentially give rise to the supporting lineage, whereas *Nr2f2*^*+*^ epithelial cells contribute to the interstitial lineage. These observations are supported by lineage-tracing experiments using *Nr2f2* and the Notch-responsive *Hes1* models, which indicate that ovarian interstitial progenitors can be identified as early as E10.5 through labeling of Notch-active and or *Nr2f2*^+^ progenitors. Nonetheless, our lineage-tracing models do not exclusively label these progenitors within the epithelium. In addition, we cannot fully exclude contributions from mesonephric progenitors, as suggested by the presence of *Gata4*^*-*^*/Gata2*^*+*^ cells in our single-cell dataset and lineage-traced cells in the mesonephros. Precise determination of progenitor origin will require strategies that exclusively label selective progenitor populations, or through applying unbiased genetic barcoding approaches combined with spatially resolved techniques.

The transcriptional heterogeneity of the coelomic epithelium in E11.5 ovary was confirmed by immunofluorescence staining of NR2F2 and NR5A1 proteins, which revealed a largely inverse expression pattern within the coelomic epithelium. This heterogeneity is observed as early as E10.0-E10.5, where NR5A1 shows mosaic expression in the coelomic epithelium (34). By E12.5, NR2F2 expression is downregulated in the ovarian epithelium (21, 35), whereas NR5A1 expression persists (36), coinciding with the contribution of the epithelium to second-wave granulosa cells based on *Lgr5* lineage tracing (12). These observations suggest that epithelial progenitors acquire molecular heterogeneity early during ovarian development; however, how *Nr2f2* and *Nr5a1* expression is initially established and dynamically regulated, and how these patterns relate to Notch signaling, requires further investigation at finer temporal resolution across early developmental stages.

We further note that, despite broad expression of Notch receptors in early gonads (20), only a subset of NR2F2^+^ cells exhibit active Notch signaling at E11.5, as indicated by both single-cell analysis and 24h short-term tamoxifen labeling. Although the fraction of labeled cells is influenced by incomplete tamoxifen labeling efficiency, the restricted pattern aligns with observations from E11.5 CBF:H2B-Venus, a tamoxifen-independent Notch transcriptional activity reporter line (20). In both cases, Notch-responsive cells are detected sporadically in the coelomic epithelium, a small number of cells within the gonad, and endothelial cells. This restricted activation may reflect local ligand availability and or concurrent expression of the Notch antagonist *Numb*. Spatially resolved approaches will be required to identify the cellular sources and contexts of Notch activation.

Importantly, our data indicate that Notch signaling is by no means merely a lineage marker, but actively drives interstitial gene program. Ectopic activation of Notch in *Nr5a1*^+^ cells promoted interstitial lineage expansion, disrupting the balance between interstitial and supporting cell populations. This result is further supported by prior studies showing that global deletion of *Hes1* reduces interstitial marker expression while increasing pre-granulosa marker expression in the ovary (19). It has also been reported that ubiquitous CreER-mediated deletion of Notch antagonist *Numb* leads to an expansion of somatic progenitor cells marked by LHX9 (20). However, while one would expect that loss of this Notch antagonist to phenocopy ectopic Notch activation, we did not observe increased expression of progenitor-associated markers including *Lhx9* and *Emx2* in our bulk RNA-seq analysis. In addition, we did not detect epithelial invagination phenotypes described in *Numb*-deleted ovaries. This suggests that Notch activation is separable from NUMB-mediated asymmetric distribution and division of the coelomic epithelium, and that loss of *Numb* alone does not necessarily result in activate Notch signaling sufficient to drive an interstitial cell transcriptional program.

Notch signaling has been shown to promote epithelial-to-mesenchymal transition in various developmental and disease contexts (37, 38). Through motif and peak-gene linkage analysis, we identified accessible chromatin regions enriched in *Nr2f2*^+^ cells containing RBPJ and/or HES1 motifs that were linked to interstitial markers, including multiple collagen genes, *Pdgfrb* and *Nr2f2* itself. These regions are also negatively associated with supporting markers such as *Wnt4* and *Kitl*. The observation that Notch-active cells are largely NR2F2^+^ and not all NR2F2^+^ cells exhibit Notch activity, suggests a potential relationship that Notch signaling may act upstream of *Nr2f2* in a context-dependent manner. Although these analyses provide predictive insight into potential regulations, they do not establish direct transcriptional causality. Definitive validation of these regulatory interactions will require chromatin immunoprecipitation-based assays or targeted perturbation of candidate regulatory elements. Moreover, supporting cells later activate Notch signaling during follicle formation without adopting mesenchymal gene expression programs (17), suggesting that Notch signaling plays context-dependent roles and that Notch activity alone is insufficient to confer mesenchymal identity, instead requiring cooperation with additional transcription factors and or signaling pathways.

Finally, our findings indicate that separation of the two somatic lineages occurs similarly through Notch pathway in both sexes. In the fetal testis, Notch signaling is activated in interstitial progenitors as early as E10.5, labeling both steroidogenic fetal Leydig cells and non-steroidogenic interstitial cells (15). However, during fetal testis development, both NR2F2 expression and Notch signaling are downregulated in a subset of interstitial cells around E13.5-E14.5 to permit fetal Leydig cell differentiation (24). Pharmacological inhibition of Notch signaling or genetic deletion of *Hes1* expands the fetal Leydig cell population, whereas sustained Notch activation maintains interstitial progenitors in an undifferentiated mesenchymal state and suppresses Leydig cell differentiation (39). In contrast, in the ovary, specification of steroidogenic theca cells occurs postnatally (14). Nonetheless, a comparable mechanism may function during theca cell differentiation, as 3βHSD^+^cells exhibit reduced NR2F2 expression (**Fig. S3B**). In summary, we propose a model in which active Notch signaling in *Nr2f2*^*+*^ progenitor cells specifies the interstitial cell lineage and separates them from the supporting cell lineage, which is characterized by absence of Notch and *Nr2f2* while positive of *Nr5a1* at the beginning of ovary formation (**Fig. 6**).

**Figure 6.**
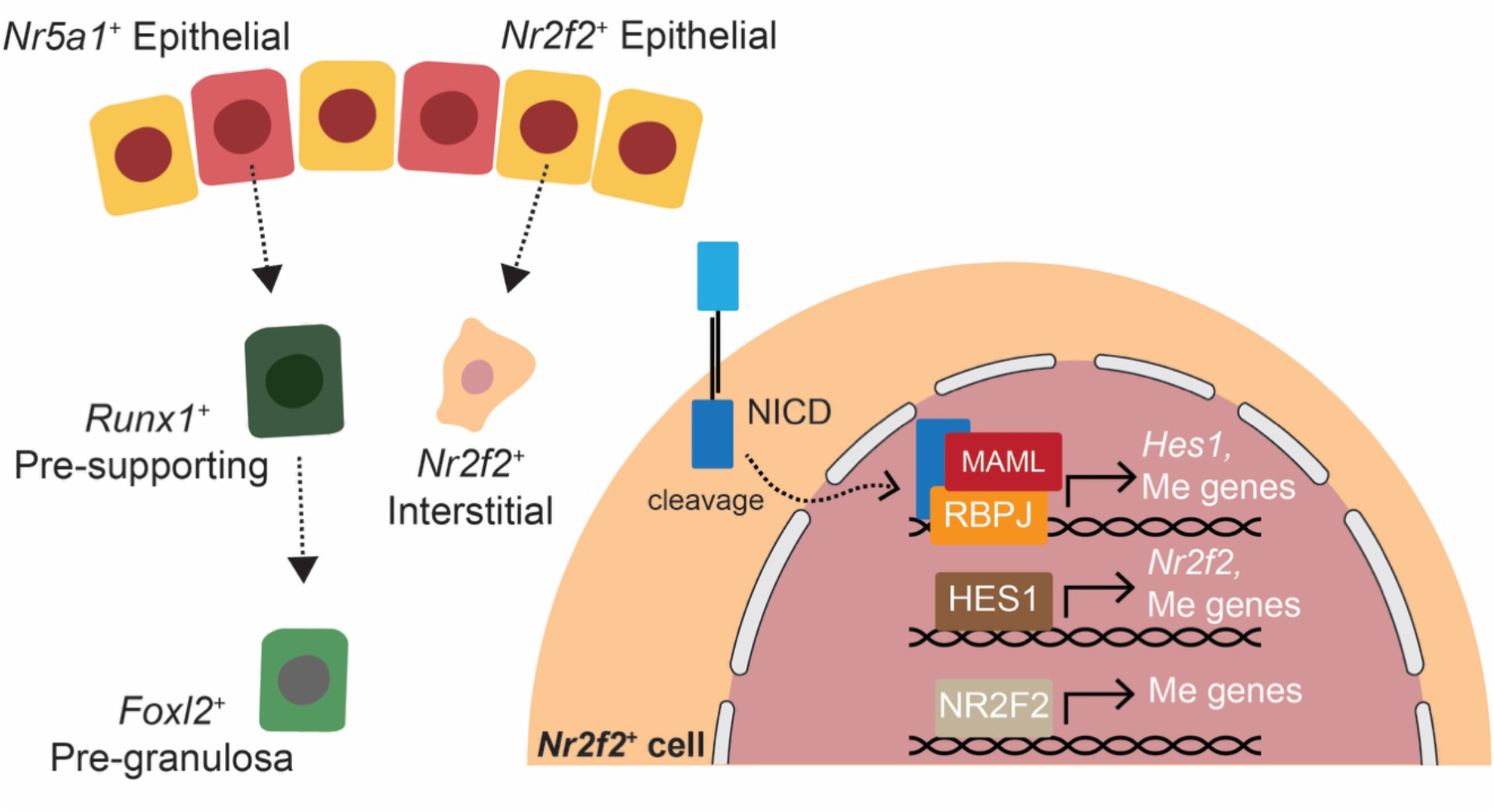
A proposed model of interstitial and supporting cell lineages differentiation during fetal ovary development. Active Notch signaling in *Nr2f2*^*+*^ progenitors specifies the interstitial lineage fate through promoting mesenchymal (Me) gene expression programs, while *Nr5a1*^+^ progenitors with low Notch activity adopt the supporting fate.

## Materials and Methods

### Animals

Animals used in this study, including C57BL/6J (JAX 000664), *Rosa*^*tdTomato9*^ (JAX 007909, B6.Cg-*Gt(ROSA)26Sor*^*tm9(CAG-tdTomato)Hze*^/J) (40), *Rosa*^*Sun1-sfGFP*^ (JAX 021039, B6;129-*Gt(ROSA)26Sor*^*tm5(CAG-Sun1/sfGFP)Nat*^/J) (41), and *Rosa*^*Notch*^ mice (JAX 008159, STOCK *Gt(ROSA)26Sor*^*tm1(Notch1)Dam*^/J) (42), were purchased from the Jackson Laboratory. *Hes1*^*CreERT2*^ (*Hes1*^*tm1(cre/ERT2)Lcm*^) mice (43) were provided by Dr. Charles Murtaugh at the University of Utah. Tg(Nr5a1-cre) (B6;D2-Tg(Nr5a1-cre)2Klp/HyJ, available at JAX 037533) mice (44) were provided by the late Dr. Keith Parker. Finally, *Nr2f2*^*CreERT2*^ mice were generated in house (35). All mice were kept in a 12-h light, 12-h dark cycle with temperature ranging 21–23°C and relative humidity ranging 40 to 50% at the animal facility at National Institute of Environmental Health Sciences (NIEHS). Animals were given chow (NIH-31M, Harlan Teklad) and water *ad libitum*. All animal studies were conducted in accordance with the NIH Guide for the Care and Use of Laboratory Animals and approved by the NIEHS Animal Care and Use Committee (Protocol 2010-0016). For time mating experiments, the day of detection of vaginal plug was considered embryonic day (E) 0.5.

### Lineage tracing

Heterozygous *Hes1*^+/CreERT2^ and *Nr2f2*^*+/CreERT2*^ male studs were crossed separately to homozygous female *Rosa*^*tdTomato9/tdTomato9*^ or *Rosa*^*Sun1-sfGFP/Sun1-sfGFP*^ mice. Once pregnant with fetuses carrying the reporter genotype of either *Hes1*^+/CreERT2^; *Rosa*^*+/tdTomato9*^, or *Hes1*^+/CreERT2^; *Rosa*^*+/Sun1-sfGFP*^, or *Nr2f2*^*+/CreERT2*^; *Rosa*^*+/tdTomato9*^, pregnant dams were dosed with 1 injection of 1mg/10g body weight tamoxifen (Sigma-Aldrich, T5648) in corn oil (Sigma-Aldrich, C8267) at E10.5, E11.5, or E13.5. Gonads and mesonephros from E11.5 female *Hes1*^+/CreERT2^; *Rosa*^*+/tdTomato9*^ embryos dosed at E10.5 were collected for histological analysis. The sex of the embryo was determined with PCR using primers that recognize the Y chromosome through Transnetyx, lnc. Ovaries from *Hes1*^+/CreERT2^; *Rosa*^*+/Sun1-sfGFP*^ fetuses dosed at E10.5 were collected at E16.5 for single-cell RNA sequencing. Whereas *Hes1*^+/CreERT2^; *Rosa*^*+/tdTomato9*^ and *Nr2f2*^*+/CreERT2*^; *Rosa*^*+/tdTomato9*^ pups were delivered through terminal C-section (45) at E18.5 or E19.5. Ovaries from F1 pups were collected at postnatal day (P) 21 for histological analysis.

### Gonadal sample collection and preparation

Gonads and mesonephros from E11.5 female B6 and lineage-traced *Hes1*^+/CreERT2^; *Rosa*^*+/tdTomato9*^ embryos were collected for whole-mount staining. Tail somite counts were recorded (19 to 21 tail somites), and the sex of the embryo was determined by PCR using Y-chromosome-specific primers through Transnetyx, lnc. Dissected gonads with attached mesonephros were fixed in 4:1 methanol (Fisher Chemical, A412): DMSO (Sigma-Aldrich, D8418) at -20°C for a minimum of 24 h prior to staining. Postnatal day 21 ovaries from lineage traced animals were fixed in 4% paraformaldehyde (PFA; Electron Microscopy Sciences, 15710) at 4°C overnight, washed in PBS, and transferred to 70% ethanol (EtOH; Sigma-Aldrich, 459844) for paraffin embedding. Ovaries from *Nr5a1*^+/Cre^; *ROSA*^*+/Notch*^ and control animals were similarly fixed in 4% PFA at 4°C overnight and washed in PBS. Embryonic day 13.5 ovaries were incubated in 30% sucrose (Sigma-Aldrich, S5016) in PBS at 4°C overnight, embedded in Tissue-Tek OCT Compound (Sakura, 4583), snap-frozen, and cut into 10 µm sections using a Leica CM3050 S cryostat before storage at - 80°C. Embryonic day 15.5 and 18.5 ovaries were transferred to 70% ethanol for paraffin embedding. Paraffin processing was performed using a Leica HistoCore PEARL processor (70% EtOH 30 min; 80% EtOH 45 min; 95% EtOH 45 min; 100% EtOH 40 min x3; xylene (J.T.Baker, 9490-03) 60, 60, and 90 min; paraffin (Leica, 39601006) 100 min x3), followed by embedding on a Leica and Arcadia embedder and sectioning at 5 µm using a Leica RM 2245 microtome.

### Immunostaining

For whole-mount staining, tissues fixed in methanol:DMSO at -20°C were washed in 50% methanol in PBS for 30 min at room temperature (RT), followed by three 1-h washes in blocking buffer (5% donkey serum (Sigma-Aldrich, S30) and 0.1% Triton X-100 (Fisher Scientific, BP151) in PBS). Tissues were incubated overnight at 4°C in blocking buffer containing primary antibodies (see **SI Table S1**). The next day, tissues were washed three times for 1 h each in blocking buffer at 4°C, followed by incubation with secondary antibodies (see **SI Table S1**) in blocking buffer at 4°C overnight. On the third day, tissues were dehydrated in sequential 1-h washes of increasing concentrations of methanol (25%, 50%, 75% with DAPI (Invitrogen D1306), and 100%) at RT, then cleared in BABB, which was made of 1:2 benzyl alcohol (Sigma-Aldrich, 402834): benzyl benzoate (Sigma-Aldrich, B6630) at 4°C for at least 24 h before imaging.

For paraffin sections, slides were deparaffinization in xylene three times for 10 min each, followed by rehydration through graded ethanol (100% EtOH 5 min x2; 95% EtOH 5 min x2; 70% EtOH 5 min x2) and a final 5-min wash in distilled water. Antigen retrieval was performed in a 1:1000 dilution of citrate acid-based antigen unmasking solution (Vector Laboratories, H-3300) by preheating the solution for 5 min at full microwave power (700 watts) and then boiling the slide for 20 min at 10% power. After cooling to RT, slides were blocked in blocking buffer for 1 h and incubated with primary antibodies overnight at 4 °C.

The following day, slides were washed three times in PBST (PBS with 0.1% Triton X-100) for 5 min each, incubated with secondary antibodies for 1 h at RT, washed three times in PBST and once in PBS, counterstained with DAPI, and mounted with ProLong Diamond Antifade Mountant (Invitrogen, P36961). Immunostaining on frozen sections was performed identically to paraffin sections but without deparaffinization, rehydration, and antigen retrieval steps. Slides were equilibrated in PBS at RT prior to blocking and antibody incubations. Imaging for all samples was performed using a Zeiss LSM 900 confocal microscope with Zen software.

### Single-cell RNA sequencing of lineage-traced cells

Ovary pairs from E16.5 *Hes1*^+/*CreERT2*^; *ROSA*^*+/Sun1-sfGFP*^ animals that were lineage-labeled at E10.5 were collected in ice-cold 1X PBS containing 0.04% bovine serum albumin (BSA; Sigma, A9576), cut into smaller pieces, and transferred into dissociation buffer: 0.04% BSA, 1.2 U/mL Dispase II (Millipore Sigma, 4942078001), 1 mg/mL collagenase B (Millipore Sigma, 11088807001), and 5 U/mL DNase I (Millipore Sigma; 101041590001) in 1X PBS. Tissue pieces were incubated at 37 °C and gently pipetted every 5 min for 20 – 30 min. To stop the enzymatic reaction, tissues were diluted with an equal volume of cold 0.04% BSA/ PBS, passed through a 40 µm FLOWMI cell strainer (SP Bel-Art, 136800040), and washed by pelleting at 350 rcf for 5 min at 4 °C, before resuspending in 110 µL cold DMEM/F12 (Thermo Fisher, 11330032). Cells were cryopreserved by adding 1:1 filtered cryoprotectant (20% DMSO in heat-inactivated FBS, HI-FBS; Thermo Fisher, 16140071) and frozen at −80°C, with each pair of the ovary in a cryovial, before long-term storage in liquid nitrogen. Once genotypes were further confirmed through PCR (Transnetyx, Inc), two cryovials containing *sfGfp*^*+*^ cells were rapidly thawed at 37°C for 1 min and pooled into one new 1.5 mL tube. This pooled sample contained cells from four ovaries from two animals carrying the *sfGfp* allele. The cryovial was rinsed three times with 300 µL of 10% HI-FBS/ DMEM/F12 pre-warmed at 37 °C using wide-bore tips, adding each rinse dropwise with gentle agitation. Cells were pelleted at 350 rcf for 5 min at 4 °C, resuspended in 50 µL 0.04% BSA/ PBS. Library preparation and sequencing were performed at the NIEHS Epigenomics and DNA Sequencing Core Facility. Cells were counted and examined for viability with AO/PI staining using a Luna FX7cell counter (Logos Biosystems). Up to 16,000 live cells were loaded into the Single Cell Chip to generate single cell emulsion in Chromium Controller with Chromium Single Cell 3’ Library & Gel Bead Kit v3.1 (10x Genomics, 1000268). Reverse transcription of mRNA and cDNA amplification were carried out following the manufacture’s instruction (10x Genomics, 1000268). The amplified cDNA was further fragmented to construct NGS libraries. Sequencing was performed on an Illumina NovaSeq 6000 using a S1 flow cell with i7 index 10bp, i5 index 10bp, achieving a depth of ∼20,000 read pairs per cell. Details of the single-cell RNA sequencing analysis are provided in the Supporting Information.

### Single-cell RNA sequencing analysis

FASTQ files were processed using *CellRanger* (v. 7.0.0) with a custom mm10 reference genome that incorporated the *sfGFP-Myc* reporter allele, which contains two copies of superfolder GFP (sfGFP) followed by six Myc epitope sequences. The reporter sequence was obtained directly from the authors (41) and is provided in the **SI Compiled Code**. *CellRanger* identified an estimated 13,994 cells with a mean of 19,890 reads per cell and a median of 1,934 detected genes per cell. The fraction of reads in cells was 79.8%, and sequencing saturation was 31.0%. Q30 scores were 95.1% (barcode), 92.7% (RNA read), and 94.9% (UMI). Mapping rates showed that 91.9% of reads aligned to the genome and 71.8% mapped confidently to the transcriptome. Single-cell data were analyzed using *Seurat* (v. 4.3.0) (46) in R (v. 4.2.1). Initial filtering included parameters of min.cells = 3 and min.features = 200. Doublets were removed using *scDblFinder* (v. 1.10.0) (47) with default settings, and ambient RNA was removed using *celda/decontX* (v. 1.12.0) (48), with the maximum iterations of the EM algorithm set to 100. The dataset was further filtered to retain cells with nFeature_RNA greater than 1,500 and less than 6,500, and percent.mt less than 25%, yielding 5,553 high-quality cells. Data were normalized and scaled, and dimensionality reduction was performed using principal component analysis (“RunPCA”) followed by UMAP embedding (“RunUMAP”, dims = 1:15). Cell clustering was performed using “FindNeighbors” (dims = 1:15) followed by “FindClusters” with a resolution of 0.1, yielding 7 transcriptionally distinct clusters. Lineage-traced cells were defined as those with “*Sfgfpmyc*” expression greater than 0 in the scaled data. Differential gene expression analysis between supporting (Pre-G/Epi and Pre-G) and interstitial clusters was performed using “FindMarkers” with default settings. Pathway enrichment analysis of DEGs (adjusted p-value < 0.05 and log2FC > 1.5) was performed using *Enrichr* (49-51).

### E11.5 XX gonadal cells joint RNA and ATAC sequencing analysis

We analyzed the previously published 10x Multiome dataset containing E11.5 XX gonadal cells (21), using *Seurat* (v. 5.2.1) (52) and *Signac* (v. 1.14.0) (53) in R (v. 4.4.1). Within the annotated E11.5 XX nuclei, we performed linear dimensional reduction using the “RunPCA” function and non-linear dimensional reduction using the “RunUMAP” function with dims = 1:30 on the scaled snRNA-seq data. Cell clustering was then performed using the “FindNeighbors” function with dims= 1:30 and the “FindClusters” function with resolution= 0.2. This generated in total 11 cell clusters. We reannotated cell clusters using markers including *Ddx4* and *Pou5f1* for primordial germ cell (PGC); *Runx1* and *Wnt4* in pre-supporting cell (PSC) (4); *Pax8* in supporting-like cells (SLC) (25); *Upk3b* and *Aldh1a*2 in the epithelial cluster (22, 23); *Ptn* and *Nr2f2* for the mesenchymal (Me) cluster; *Hba-a1* in the erythrocyte (Ery) cluster; *Cx3cr1* and *Fcgr1* in the immune cluster; endothelial cluster is positive of *Pecam1* but negative of PGC gene expression; mesonephric tubule (MT) is positive of *Pax2*; and finally, *Eya1* and *Phox2b* for adrenosympathetic (AS) cells (2, 25, 54, 55).

For ATAC analysis, we applied the peak set that was called on individual RNA clusters from E11.5-E13.5 dataset. Within the annotated E11.5 XX nuclei, we normalized the data using latent semantic indexing (LSI) by applying “RunTFIDF”, “FindTopFeatures”, and “RunSVD” functions in *Signac*. Visualization was performed using UMAP with dims = 2:30, reduction = “lsi”. ATAC clustering was performed using the “FindNeighbors” function with dims= 2:30 and the “FindClusters” function with resolution= 0.3. This generated in total 9 clusters. Joint UMAP was generated with the “FindNultiModalNeighbors” with dims = 1:30 for “pca” and 2:30 for “lsi”. Peak gene linkage was analyzed using the “LinkPeaks” function in *Signac* on the entire E11.5 XX dataset. Velocity inference was performed using *MultiVelo* (v. 0.1.2) (26) in Python (v. 3.9.23). Joint UMAP coordinates generated in *Seurat* were supplied as the low-dimensional embedding for model initialization. Chromatin accessibility input was derived from peak-associated gene activity scores computed using the “GeneActivity” function in *Signac*. All remaining parameters were run using the default *MultiVelo* settings.

Differentially expressed genes (DEG) between the two epithelial clusters were determined using the “FindMarkers” function in *Seurat* with default settings. Pathway analysis on DEGs with adjusted p-value < 0.05 and log2FC >1 was performed using *Enrichr* (49-51). Differentially accessible peaks (DAP) between the two epithelial clusters were determined using the “FindMarkers” function based on the LR test with min.pct = 0.01 and logfc.threshold = 0.1.

Motif enrichment was performed using the “FindMotifs” function in *Signac*. Motif position frequency matrices (PFM) for vertebrates were obtained from the 2022 JASPAR Core database (56) with in total 1,956 elements. Background peaks were selected to match the GC content in the peak set by using the “AccessiblePeaks,” “GetAssayData”, and “MatchRegionStats” functions in *Signac*. Enriched motifs were filtered with p.adjust < 0.05 and fold.enrichment > 1.25. *MotifScan* (v. 1.3.0) (57) was used to determine the genomic position of linked peaks to DEGs to identify predicted target genes of transcription factors.

### Bulk RNA-sequencing of Notch-induced ovaries

Embryonic day 13.5 Notch-induced (Tg(Nr5a1-cre)+; *ROSA*^*+/Notch*^) and control (Tg(Nr5a1-cre)+;; *ROSA*^+/+^) ovary pairs were collected for bulk RNA-sequencing. As matings were performed by crossing either male or female Tg(Nr5a1-cre) with *ROSA*^*+/Notch*^, we collected three pairs of Notch-induced ovaries from *ROSA*^*+/Notch*^ dams, three pairs of Notch-induced ovaries from Tg(Nr5a1-cre) dams, three pairs of control ovaries from *ROSA*^*+/Notch*^ dams, and three pairs of control ovaries from Tg(Nr5a1-cre) dams, yielding six biological replicates for each genotype. Each ovary pair was dissected and snap-frozen in 50 µL of extraction buffer from the Arcturus PicoPure RNA Isolation Kit (Applied Biosystems, 12204-01) and stored at −80°C. For RNA isolation, samples were thawed on ice, vortexed, mixed with an equal volume of 70% ethanol, and applied to pre-conditioned columns. On-column DNA digestion was performed using the RNase-Free DNase Set (Qiagen, 79254), at a final concentration of around 0.34 U/µL for 15 min at RT. After two washes, RNA was eluted in 11 µL of elution buffer. RNA quantity and quality were measured using the Qubit RNA High Sensitivity kit (Invitrogen, Q32852) and the High Sensitivity RNA Screen Tape Analysis (Agilent, 5067-5579) on the Agilent TapeStation 4200.

Library preparation and sequencing were performed at the NIEHS Epigenomics and DNA Sequencing Core Facility. Libraries were made using the TruSeq Stranded mRNA (Illumina, 20020594) according to the manufacturer’s protocol. Approximately 250 ng of total RNA from each sample was used for poly(A) mRNA enrichment followed by cDNA synthesis and library construction. Sequencing was performed on an Illumina NextSeq 500 sequencer using a SP flow cell to generate paired-end 150-bp reads. Sequencing yielded ∼80 million reads per sample. FASTQ files were generated using bcl2fastq/DRAGEN with default parameters. Details of the bulk RNA sequencing analysis are provided in the Supporting Information.

### Notch-induced ovaries analysis

Raw paired-end FASTQ files were trimmed using *Trim Galore* (v. 0.6.10) (58), with the following parameters: --quality 20, --illumina, --stringency 5, --length 20, and --paired. The mm10 reference genome and NCBI RefSeq GTF annotation were downloaded from the UCSC Genome Browser on January 9, 2025 and used to generate a *STAR* (v. 2.6.0c) (59) genome index. Trimmed reads were aligned to the mm10 reference using STAR, producing coordinate-sorted BAM files, which were indexed with *samtools* (v. 1.18). Gene-level quantification was performed with featureCounts (Subread, v. 2.0.6) (60) using the RefSeq annotation, specifying paired-end mode (-p) and reverse-stranded orientation (-s 2). Differential gene expression analysis was conducted using the Wald test as implemented in *DESeq2* (v. 1.44.0) (61). Genes with an adjusted *p*-value < 0.05 (Benjamini–Hochberg correction) were considered significant. Pathway analysis on DEGs with adjusted p-value < 0.05 and log2FC >1 was performed using *Enrichr* (49-51). Single-cell deconvolution was performed using *MuSiC* (v. 1.0.0) (62) with E13.5 XX ovarian cells from the 10X Multiome dataset served as our single-cell reference.

### Statistical analyses

Differentially expressed gene (DEG) analysis for single-cell and single-nucleus RNA-seq datasets was performed using the Wilcoxon Rank Sum test as implemented in *Seurat*, with min.pct = 0.01 and logfc.threshold = 0.1. Genes with an adjusted *p*-value < 0.05 were considered significantly differentially expressed. DEG analysis for bulk RNA-seq was conducted using *DESeq2* with standard normalization and dispersion estimation procedures using default parameters. Differentially accessible peaks (DAPs) from snATAC-seq data were identified using the likelihood ratio (LR) test implemented in *Signac*, with min.pct = 0.01, logfc.threshold = 0.1, and adjusted *p*-value < 0.05 considered significant. Differences in *MuSiC*-estimated cell-type proportions between groups were assessed using the Wilcoxon Rank Sum test performed separately for each cell type. Resulting *p*-values were corrected for multiple testing using the Benjamini–Hochberg false discovery rate (FDR) procedure.

## Data availability

Analyses using E11.5 and E13.5 XX gonads were performed using sequencing data in the Gene Expression Omnibus (GEO) under accession GSE288206 (21). Newly sequenced data, including lineage-traced E16.5 ovarian single-cell RNA-seq and E13.5 Notch-induced ovarian bulk RNA-seq will be deposited upon publication. The code used for all analyses is included in the Supporting Information. All remaining data are available in the main text and Supporting Information.

## Acknowledgments

We thank the NIEHS Comparative Medicine Branch for mouse colony maintenance, as well as the Epigenomics and DNA Sequencing Core and the Integrative Bioinformatics Support Group, especially B. Papas, Y. Zhang, M. Cook, J. Malphurs, and F. Day for their assistance with sequencing and data analysis. We are grateful to current and past members of the Yao lab, especially K. Rodriguez and B. Nicol for their valuable input and assistance with experiments. We also thank B. Capel and L. Mork for helpful discussions and feedback. This work was supported by the Intramural Research Program ZIAS102965 of the NIH, National Institute of Environmental Health Sciences to H.H.-C.Y. and the Lalor Foundation Postdoctoral Fellowship to Y.-Y.C.

