## Supplemental Information for "Specification of the interstitial cell lineage in the fetal mouse ovary requires balanced Notch signaling"

### Supplementary Figures

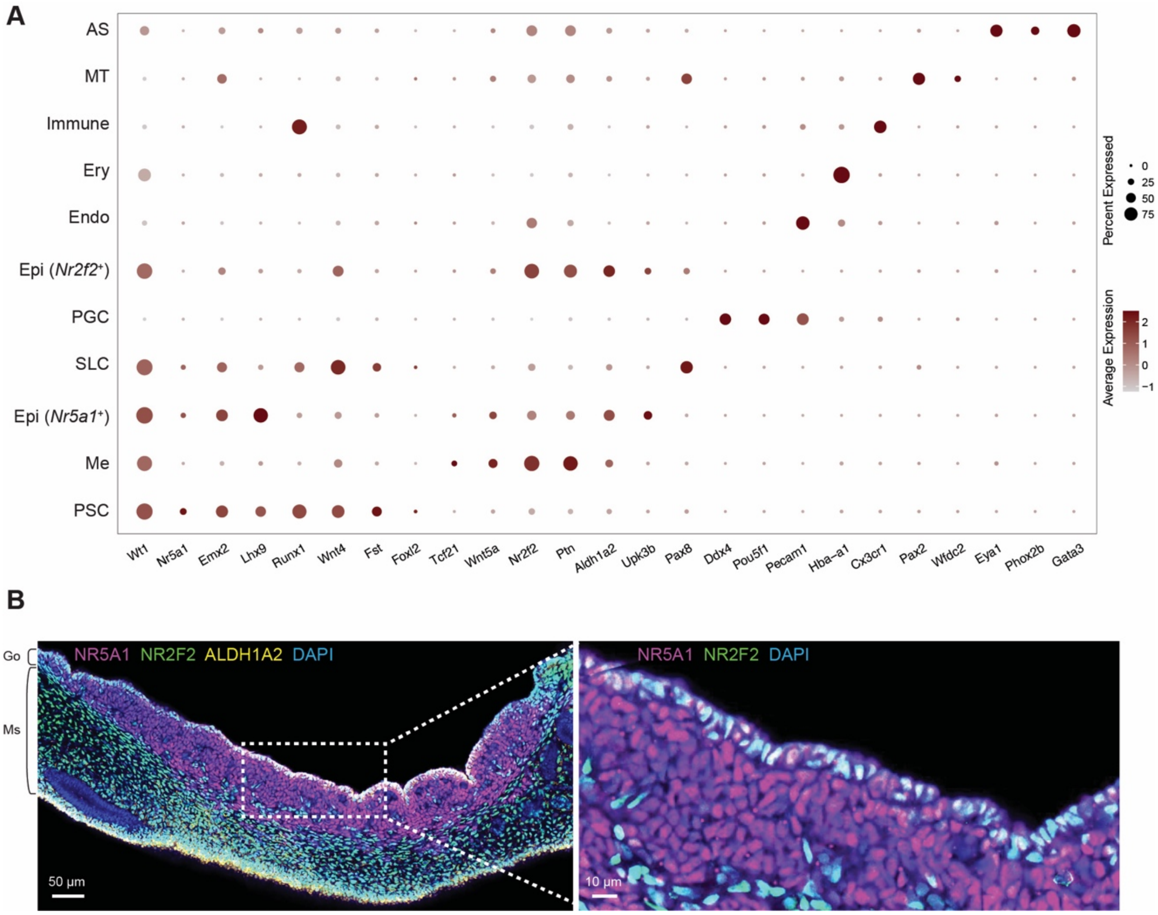


**Fig. S1. (A)** Dot plot of marker gene expression in all annotated clusters in E11.5 XX gonads. **(B)** Immunostaining of E11.5 (tail somite 21) wildtype XX gonad (Go) and mesonephros (Ms) with NR5A1 (magenta), NR2F2 (green), ALDH1A2 (yellow), and DAPI (blue). Scale bar= 50µm (left) and 10µm (inset).

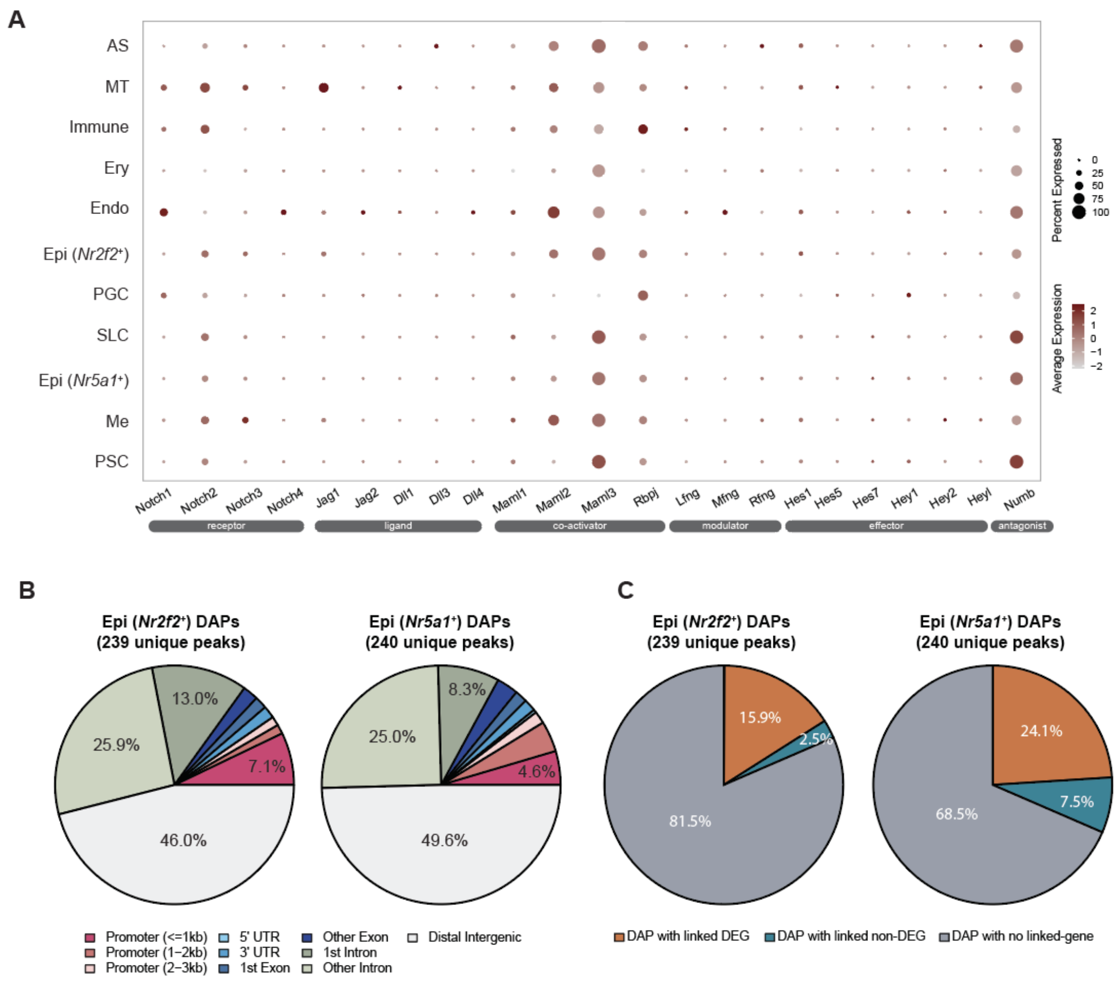


**Fig. S2. (A)** Dot plot of all Notch components expression. **(B-C)** Peak annotation of genomic location (B) and peak-gene linkage (C) of *Nr2f2*$^+$ and *Nr5a1*$^+$ epithelial enriched differentially accessible peaks (DAPs), totaling 239 and 240 unique peaks, respectively. We categorized DAPs into 1) positively or negatively linked to any DEGs, 2) positively or negatively linked to genes that were not DEGs between the two epithelial clusters, or 3) not linked to any genes at all. For the first category, we defined DAPs with at least one linked DEG as “DAP with linked DEG”. In the second category, when a DAP was linked to a gene, however the gene itself was not a DEG between the two epithelial clusters, we classified it “DAP with linked non-DEG". In the third category, when a DAP was not associated with any genes within the TSS ± 500kb linkage distance across cell types it was assigned as “DAP with no linked-gene”.

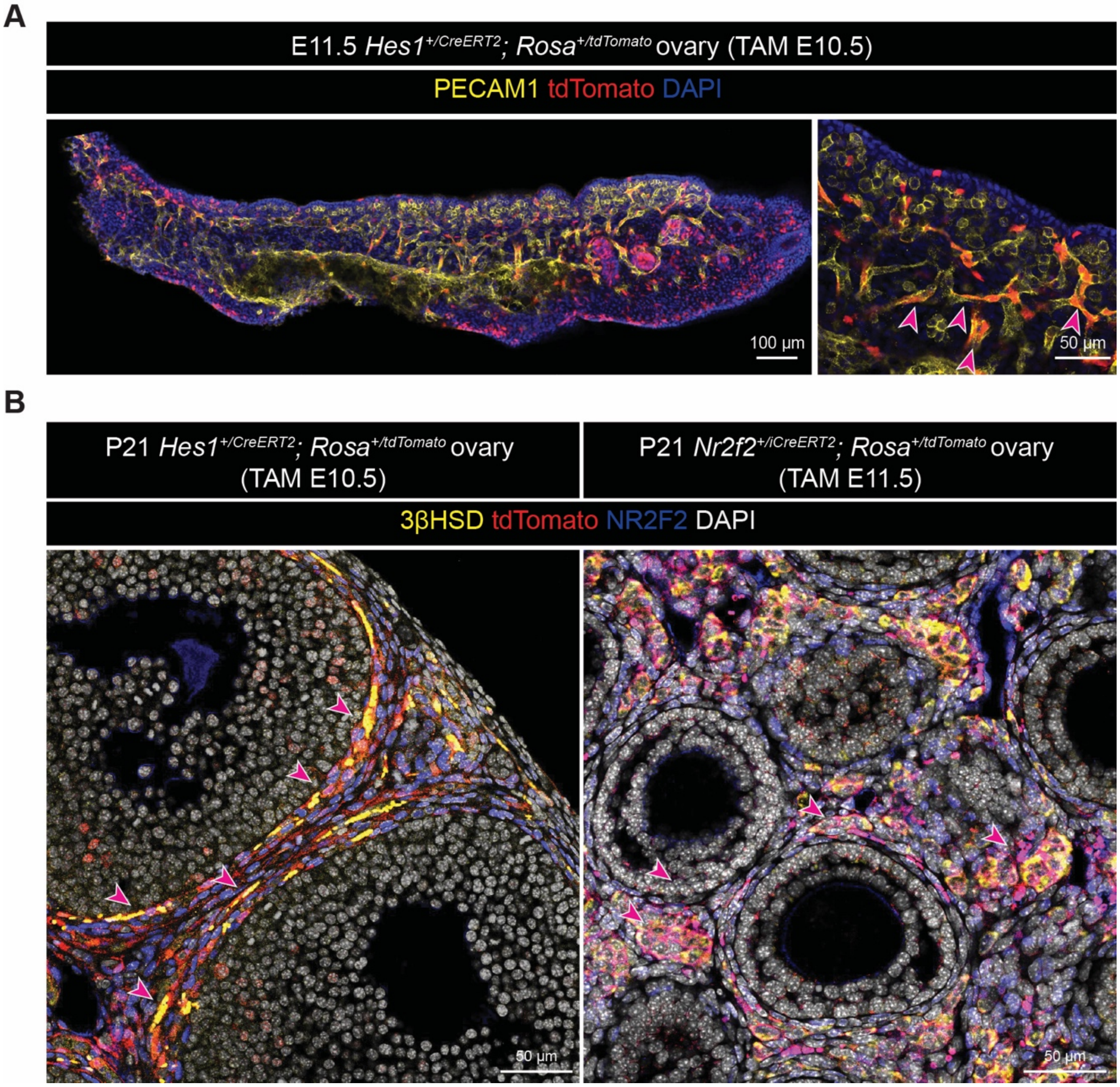


**Fig. S3. (A)** Immunostaining of E11.5 (tail somite 19) *Hes1*$^{+/CreERT2}$; *ROSA*$^{+/Sun1\text{-}sfGFP}$ ovaries lineage-labeled at E10.5, marked with PECAM1 (yellow), tdTomato (lineage-traced cells, red), and DAPI (blue). Red arrows mark lineage-traced cells expressing PECAM1. Scale bar= 100µm (left) and 50µm (inset, right). **(B)** Immunostaining of P21 *Hes1*$^{+/CreERT2}$; *ROSA*$^{+/tdTomato}$ (left) and *Nr2f2*$^{+/iCreERT2}$; *ROSA*$^{+/tdTomato}$ (right) ovaries lineage-traced at E10.5 (left) or E11.5 (right), marked with 3βHSD (yellow), tdTomato (lineage-traced cells, red), NR2F2 (blue), and DAPI (white). Red arrows mark lineage-trace cells expressing 3βHSD. Scale bar= 50µm.

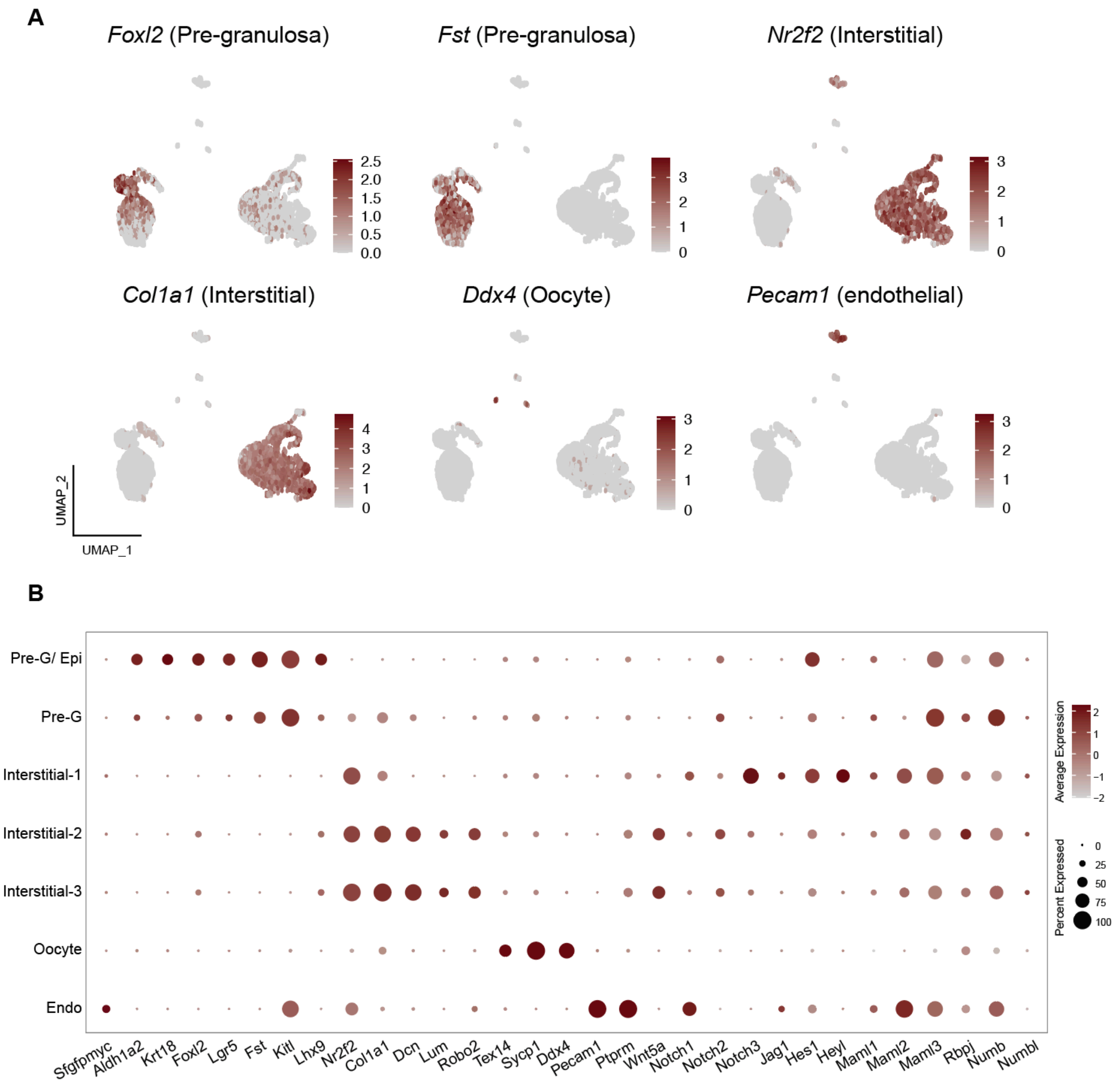


**Fig. S4. (A)** Feature plots of selected marker gene expression in E16.5 *Hes1*$^{+/CreERT2}$; *ROSA*$^{+/Sun1\text{-}sfGFP}$ ovaries lineage-labeled at E10.5. **(B)** Dot plot of the expression of marker genes and Notch signaling components across identified cell types.

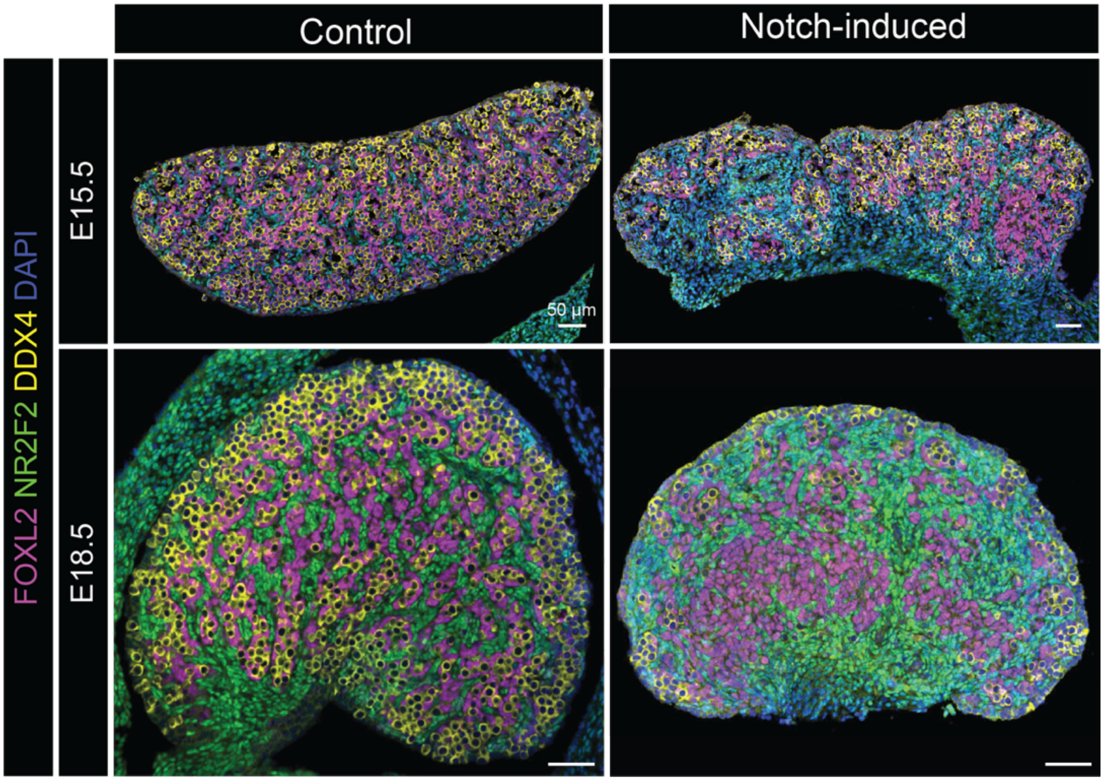


**Fig. S5.** Immunofluorescence of control and Notch-induced ovaries at E15.5 (top) and E18.5 (bottom), marked with FOXL2 (magenta), NR2F2 (green), DDX4 (yellow), and DAPI (blue). Scale bar= 50µm.

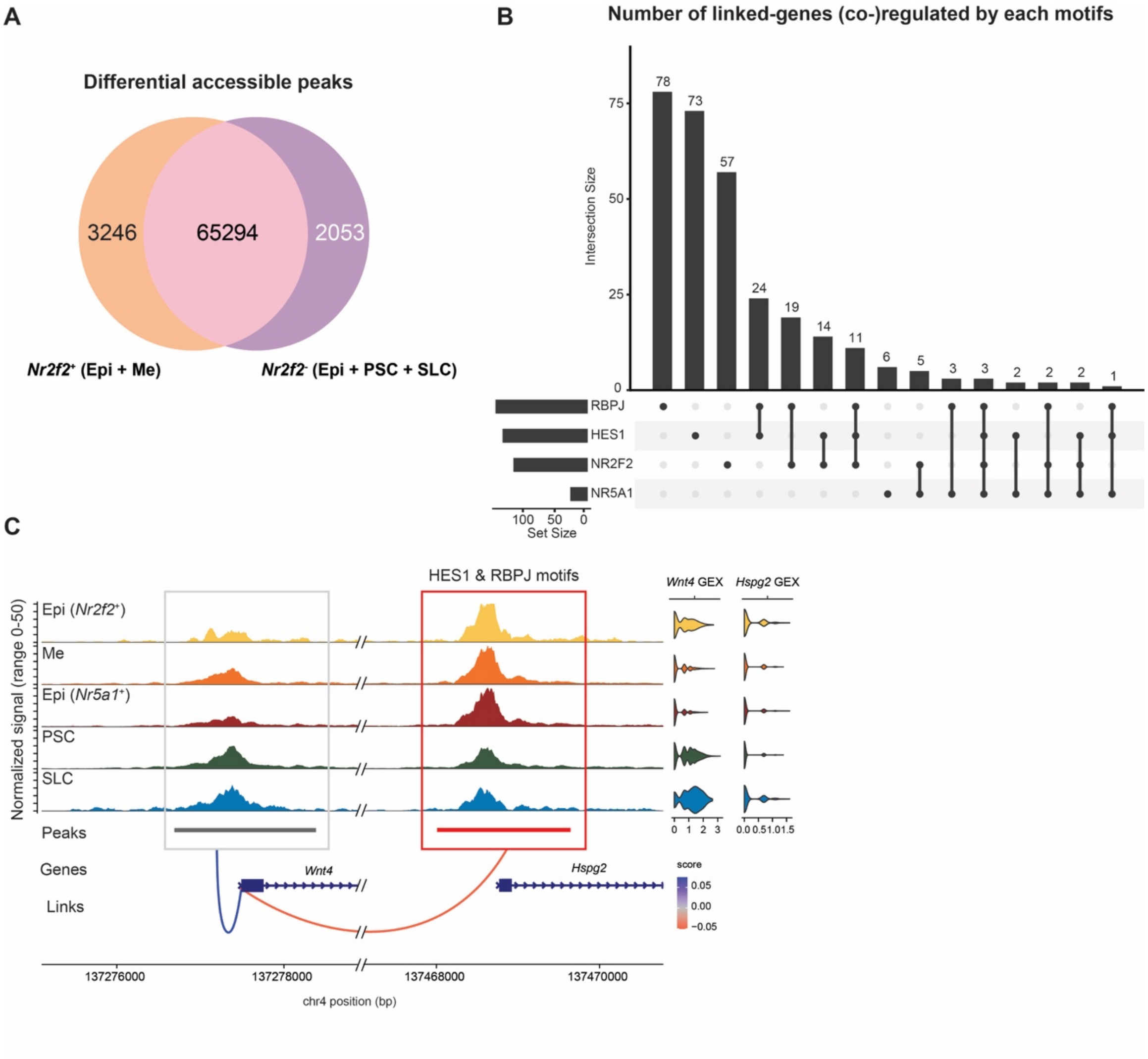


**Fig. S6. (A)** Venn diagram illustrating the number of enriched DAPs obtained by comparing all *Nr2f2*$^+$ cells (*Nr2f2*$^+$ epithelial and mesenchymal) versus *Nr2f2*$^-$ cells (*Nr5a1*$^+$ epithelial, pre-supporting, and supporting-like cells). **(B)** Upset plot summarizing positively linked DEGs identified in Fig. 5B and their association with peaks containing RBPJ, HES1, NR2F2, or NR5A1 motifs. The upper panel shows the number of genes linked to each motif combination, and the lower-left panel shows the size of each motif-defined set. **(C)** Coverage plots of the *Wnt4* and *Hspg2* loci. *Wnt4* expression was negatively associated with the *Hspg2* promoter peak that harbor both HES1 and RBPJ motifs but showed no positive association with *Hspg2* expression itself. Despite positively linked to its gene expression, the *Wnt4* promoter peak itself was not differentially accessible between *Nr2f2*$^+$ and *Nr2f2*$^-$ populations and did not contain Notch-related motifs.

**Tables**

*Primary antibodies*

| | | | | |
|---|---|---|---|---|
| NR2F2 (COUP-TFII) | Mouse | R&D Systems | PP-H7147-00 | 1:200 |
| NR5A1 (SF1) | Rat | Cosmo Bio | KO610 | 1:200 |
| ALDH1A2 | Rabbit | Sigma-Aldrich | HPA010022 | 1:200 |
| tdTomato (Living Colors® DsRed) | Rabbit | Takara | 632496 | 1:1000 |
| tdTomato (RFP) | Chicken | Rockland | 600-901-379 | 1:100 |
| FOXL2 | Goat | Novus | NB100-1277 | 1:200 |
| CYP17A1 | Goat | Santa Cruz | sc-46081 | 1:100 |
| 3βHSD | Rabbit | Cosmo Bio | K0607 | 1:500 |
| PECAM1 (CD31) | Rat | BD Biosciences | 550274 | 1:500 |
| TRA98 (GCNA) | Rat | DiagnoCine | 73-003 | 1:1000 |
| DDX4 | Rabbit | Abcam | Ab13840 | 1:400 |
| Laminin | Rabbit | Sigma-Aldrich | L9393 | 1:200 |

*Secondary antibodies*

| | | | |
|---|---|---|---|
| Alexa Fluor 488 donkey anti-mouse IgG | Invitrogen | A21202 | 1:200 |
| DyLight 650 donkey anti-rat IgG | Invitrogen | SA5-10029 | 1:200 |
| Alexa Fluor 568 donkey anti-rabbit IgG | Invitrogen | A10042 | 1:200 |
| Alexa Fluor 647 donkey anti-goat IgG | Invitrogen | A21447 | 1:200 |
| Cy™3 AffiniPure® donkey anti-chicken IgY (IgG) | Jackson Immuno Research | 703-165-155 | 1:200 |

**Table S1.** Antibody list
